# Ultrasonic potentiation of ketamine neuromodulation

**DOI:** 10.64898/2026.07.29.741494

**Authors:** Kanchan Sinha Roy, Payton Martinez, Sedona N. Ewbank, Kenneth Shinozuka, Mahaveer Purohit, Yun Xiang, Raag D. Airan

**Author notes:** These authors contributed equally (Kanchan Sinha Roy, Payton Martinez).

## Abstract

The psychiatric utility of ketamine is limited by its dissociative and systemic side effects. Recently, to enable precision ketamine pharmacotherapy, we introduced SonoKet, ketamine-loaded acoustically activatable liposomes that enable focused ultrasound (FUS)-targeted ketamine delivery to millimeter-sized brain regions. In initial studies, we observed that SonoKet uncaging targeted ketamine to the ultrasound-treated brain region, while inducing greater electrophysiologic and behavioral functional effects than dose-matched free ketamine. To further define these uncaging-potentiated neuromodulatory effects, we used solid-phase microextraction (SPME) coupled to LC-MS/MS to investigate the effect of ultrasound and SonoKet uncaging on key neurotransmitters in real-time. SPME probes were used to sample ketamine, its metabolites, and glutamate, GABA, serotonin (5-HT), and dopamine in the medial prefrontal cortex (mPFC), nucleus accumbens (NAc), and retrosplenial cortex (RsC) of awake rats. Sampling occurred before and after intravenous administration of either SonoKet, free ketamine, or saline, with FUS targeted to either a frontolimbic or caudal brain region. FUS alone did not yield significant changes in neurotransmitter concentration, nor did it affect the pharmacodistribution of free ketamine. In contrast, FUS generally increased the neurotransmitter response to free ketamine, suggesting an ultrasonic potentiation of ketamine neuromodulation. SonoKet (0.75 mg/kg) uncaging with FUS elicited further elevations in glutamate, GABA, and 5-HT within the FUS-targeted region, along with an increase in dopamine in the NAc when the frontolimbic region was sonicated. These increases were similar to or higher than those induced by 10 mg/kg free ketamine alone or 0.75 mg/kg free ketamine combined with FUS, especially with frontolimbic SonoKet uncaging. Altogether, FUS potentiates ketamine-induced neuromodulation, with spatially specific and synergistically greater effects when ketamine is spatially localized via ultrasonic uncaging. This strategy could augment ketamine pharmacotherapy for psychiatric diseases, while limiting its dissociative and abuse liabilities.

**Highlights:**

- Focused ultrasound potentiated ketamine-driven glutamate, serotonin, and dopamine release
- Localized ketamine delivery with SonoKet uncaging drove synergistically greater region-specific neurochemical responses
- Uncaging boosts ketamine effects at a fraction of the ketamine dose

**Graphical Abstract:** 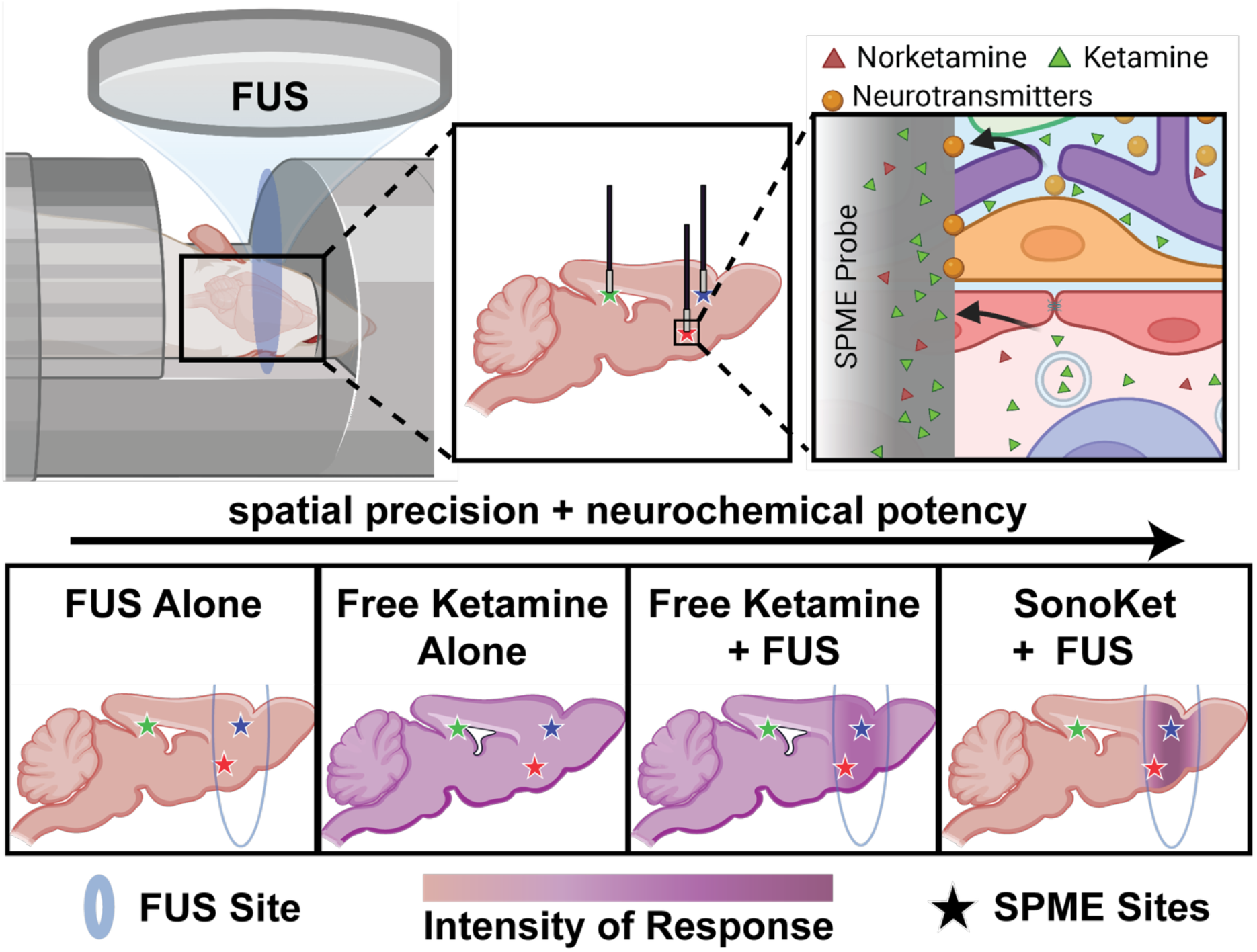

## Introduction

Transcranial focused ultrasound (FUS) is a noninvasive neuromodulation modality that can target deep brain structures with millimeter-scale precision [1–4]. FUS can also facilitate targeted drug delivery, including via on-demand drug release from carrier particles [1–7]. Its deep penetration and high spatial resolution compared to other noninvasive neuromodulation modalities make FUS a compelling platform in both research and therapeutic settings [8–10].

Currently, pharmacology remains the dominant form of neuropsychiatric treatment. However, pharmacologic therapies tend to have significant side effects that limit tolerability and compliance. For instance, ketamine, a dissociative anesthetic and recreational drug, is also an approved rapid-acting antidepressant [11–14]. However, the dissociative, hallucinogenic, abusable, addictive, and systemic side effects of ketamine limit its tolerability and adoption by both psychiatric providers and patients [15–17]. These limitations arise partly because conventional systemic dosing delivers ketamine to both therapeutic and off-target neural circuits, producing widespread neurochemical effects across the brain, in addition to off-target action in the rest of the body. Importantly, ketamine’s beneficial and adverse actions may be separable at the circuit level. The medial prefrontal cortex (mPFC) underpins the antidepressant and affective actions of ketamine [18,19], whereas the retrosplenial cortex (RsC) is linked to its dissociative effects [20]. In parallel, the nucleus accumbens (NAc), a key hub of mesolimbic dopaminergic signaling, may underlie ketamine’s psychostimulant properties, abuse liability, and other adverse neurobehavioral outcomes [21–24]. Accordingly, there is a strong interest in strategies that can maximize the beneficial neuroplastic and therapeutic effects of drugs like ketamine while minimizing their systemic and nonspecific effects via circuit-specific delivery.

Toward this goal, we recently developed SonoKet – acoustically activatable liposomes (AALs) loaded with ketamine that release encapsulated ketamine upon FUS stimulation, enabling on-demand “uncaging” within a targeted brain region [25]. Our prior work demonstrated that SonoKet uncaging locally releases ketamine at ultrasound-targeted brain regions, limits exposure of off-target brain and bodily regions, and produces region-specific electrophysiological and behavioral effects that are each greater than dose-matched unencapsulated ketamine [25]. Specifically, uncaging of SonoKet at a frontolimbic region containing the mPFC was associated with electrophysiologic and behavioral signatures consistent with antidepressant-like activity, whereas targeted release at a caudal region containing the RsC produced distinct electrophysiological responses associated with ketamine-induced dissociation [25]. Each of these effects were to a greater extent than dose-matched systemic free ketamine.

While these results show that SonoKet uncaging potentiates the effects of ketamine, it is unclear how SonoKet uncaging yields this potentiation, given that the local ketamine concentration achieved with uncaging was only marginally greater than that achieved with dose-matched free ketamine. Does FUS function only as a physical trigger for ketamine release from SonoKet, or does FUS also potentiate ketamine-induced neural responses? This question may be elucidated by direct measurement of ketamine, its metabolites, and endogenous neurotransmitters within specific brain regions following uncaging. However, real-time monitoring of neurochemical dynamics within discrete brain regions has been technically challenging. Recently, solid-phase microextraction coupled with mass spectrometry (SPME-MS) has emerged as a powerful *in vivo* chemical biopsy approach for neurochemical monitoring, using minimally invasive microprobes that function as localized sampling probes, passively sampling extracellular concentrations of numerous metabolites and neurotransmitters in a multiplexed manner, enabling repeated sampling with minimal tissue disruption and parallel measurements across multiple regions [26–34]. Unlike conventional approaches that require larger sampling volumes or continuous perfusion, SPME relies on diffusion-based, non-exhaustive extraction onto a biocompatible coating [35–37]. In awake non-human primates, SPME enabled reliable detection of pharmacology-driven neurochemical changes across cortical and striatal circuits with minute-scale temporal resolution [38,39].

In this study, we combined ultrasonic ketamine uncaging with *in vivo* SPME sampling coupled with LC-MS/MS to monitor ketamine, its metabolites, and key neurotransmitters over time in awake rodents. We tested whether SonoKet uncaging at either the same frontolimbic region (containing the mPFC and NAc) or caudal region (containing the RsC) as in our prior study generates distinct neurochemical signatures compared with administration of free, unencapsulated ketamine, given the distinct electrophysiologic and behavioral effects of uncaging in each region in our prior study [25]. We quantified time-resolved extracellular levels of glutamate, GABA, dopamine, and serotonin, along with ketamine and its key metabolites, across the mPFC, NAc, and RsC following saline, free ketamine (0.75 mg kg⁻¹ or 10 mg kg⁻¹), or SonoKet (0.75 mg kg⁻¹) administration with or without FUS targeted to either frontolimbic or caudal region. By systematically comparing FUS treatment alone, systemic free ketamine combined with FUS, and spatially localized ketamine delivery through SonoKet uncaging, we tested whether ultrasound and ketamine act independently or synergistically to reshape neurotransmitter dynamics across functionally distinct brain circuits. These findings provide mechanistic insight into the neurochemical basis of therapeutic versus dissociative actions. They further establish a translational framework for maximizing the therapeutic potential and minimizing the side effect profile of drugs like ketamine via ultrasonic potentiation, extending our previously reported platform for ultrasonic drug uncaging [25].

## Results

### SonoKet uncaging targets ketamine to the sonicated region

We used LC-MS/MS analysis to compare the concentrations of ketamine, its metabolites, and key neurotransmitters extracted via SPME from the mPFC, NAc, and RsC following free ketamine or SonoKet administration with or without ultrasound application (Fig. 1b-e). Burr hole craniectomies (∼1 mm diameter) were performed the day before the experiment to facilitate brain SPME probe placement. The ultrasound transducer lateral full-width at half-maximum (FWHM) is ∼6.25 mm and the axial FWHM is ∼22 mm (Supplementary Fig. 1), covering the full dorsal– ventral axis of the rat brain and allowing control sampling only lateral or anteroposterior to the focus rather than axially/dorsoventrally. Two baseline SPME samples were collected before treatment, followed by three post-treatment samples at 7–12 min, 30–35 min, and 60–65 min from the start of the 5 min treatment period (i.e., 2-7 min, 25-30 min, and 55-60 min respectively from the end of the 5 min treatment period) (Fig. 1a). Awake rats received intravenous (IV) infusion of saline vehicle, free ketamine HCl at 0.75 mg kg⁻¹ with or without FUS, free ketamine HCl at 10 mg kg⁻¹ without FUS, or SonoKet at 0.75 mg kg⁻¹ with or without FUS (n = 4 animals per group). Ultrasound was applied during the final 2.5 min of the 5 min IV infusion following the same protocol as used in our prior study [25]. Following cessation of infusion and sonication, SPME sampling was performed simultaneously in the mPFC, NAc, and RsC. This design enabled direct comparison of five mechanistically distinct conditions: FUS alone, free ketamine HCl with or without FUS, SonoKet infusion alone and spatially-localized ketamine release through SonoKet uncaging.

**Figure 1:**
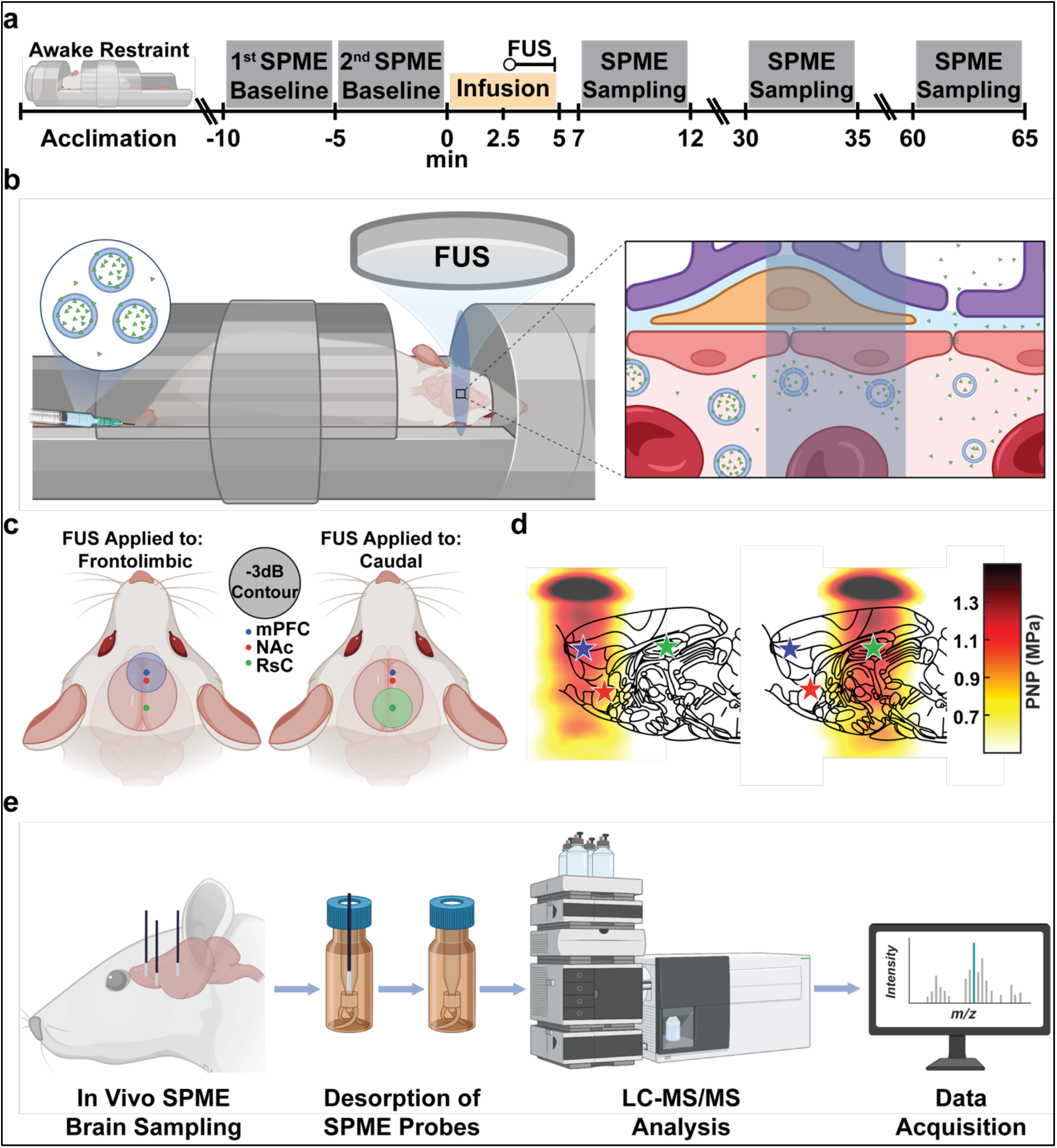
Experimental overview, focused ultrasound (FUS) targeting, awake restraint acclimation, and solid-phase microextraction (SPME) coupled LC–MS/MS analysis. (a) Experimental timeline, including animal acclimation to awake restraint, treatment, and SPME sampling periods. Rats were acclimated to restraint prior to two baseline (t = -10.0 to -5.0 and t = -5.0 to -0.0 min) SPME samplings (t = 2.5–5.0 min, followed by IV infusion of saline, ketamine HCl or SonoKet (t = 0.0–5.0 min) with or without sonication (t = 2.5–5.0 min) and subsequent SPME sampling at defined time points: 7-12 min, 30-35 min and 60-65 min. (b) Schematic of tail vein IV infusion of saline, ketamine HCl, or SonoKet administration under awake restraint animals and transcranial FUS application to enhance drug penetration and induce neuromodulation. (c) FUS targeting scheme for the frontolimbic (left, blue circle) and caudal (right, green circle) brain regions, with overlaid −3 dB pressure contours (d) simulated ultrasound beam profiles overlaid on sagittal brain schematics [67] with two targeting regions, and three SPME sampling sites with blue (mPFC), red (NAc), and green (RsC) asterisk signs. (d) Workflow for SPME-based real-time neurochemical monitoring workflow, including *in vivo* SPME sampling, sample processing, LC–MS/MS analysis, and data acquisition.

We first examined whether FUS altered the regional pharmacodistribution of free ketamine HCl or SonoKet-derived ketamine across the mPFC, NAc, and RsC (Fig. 2). Following IV infusion of 0.75 mg kg⁻¹ free ketamine HCl, SPME sampling showed that ketamine increased rapidly in all sampled regions during the first post-treatment window (7–12 min from infusion start) and then progressively declined over time (Fig. 2b, top row). FUS applied to a frontolimbic region containing the mPFC and NAc did not significantly alter free ketamine exposure: ketamine levels in the mPFC, NAc, and RsC were similar across sonicated and unsonicated conditions. Norketamine was detected at lower levels than ketamine and showed a delayed, lower-amplitude profile (Fig. 2b, bottom row). These data indicate that FUS did not measurably alter the pharmacodistribution of free ketamine HCl or its metabolites. Ketamine exposure increased markedly across all regions following IV infusion of 10 mg kg⁻¹ free ketamine HCl without ultrasound; nonsignificant trends towards regional differences in peak uptake seen at this higher dose may represent differential blood flow or receptor profiles of each region. The highest concentrations were again detected during the initial 7–12 min sampling window, followed by progressive clearance at later times (Fig. 2b, top row). Norketamine levels also increased after 10 mg kg⁻¹ free ketamine HCl and remained detectable across all sampling windows (Fig. 2b, bottom row). Hydroxynorketamine was largely undetectable after 0.75 mg kg⁻¹ free ketamine with or without FUS but became measurable after 10 mg kg⁻¹ ketamine HCl, consistent with dose-dependent systemic metabolism (Supplementary Fig. 3, top row). This dose-dependent profile confirms that the SPME coupled with LC-MS/MS captured pharmacologically relevant changes in regional ketamine exposure.

**Figure 2:**
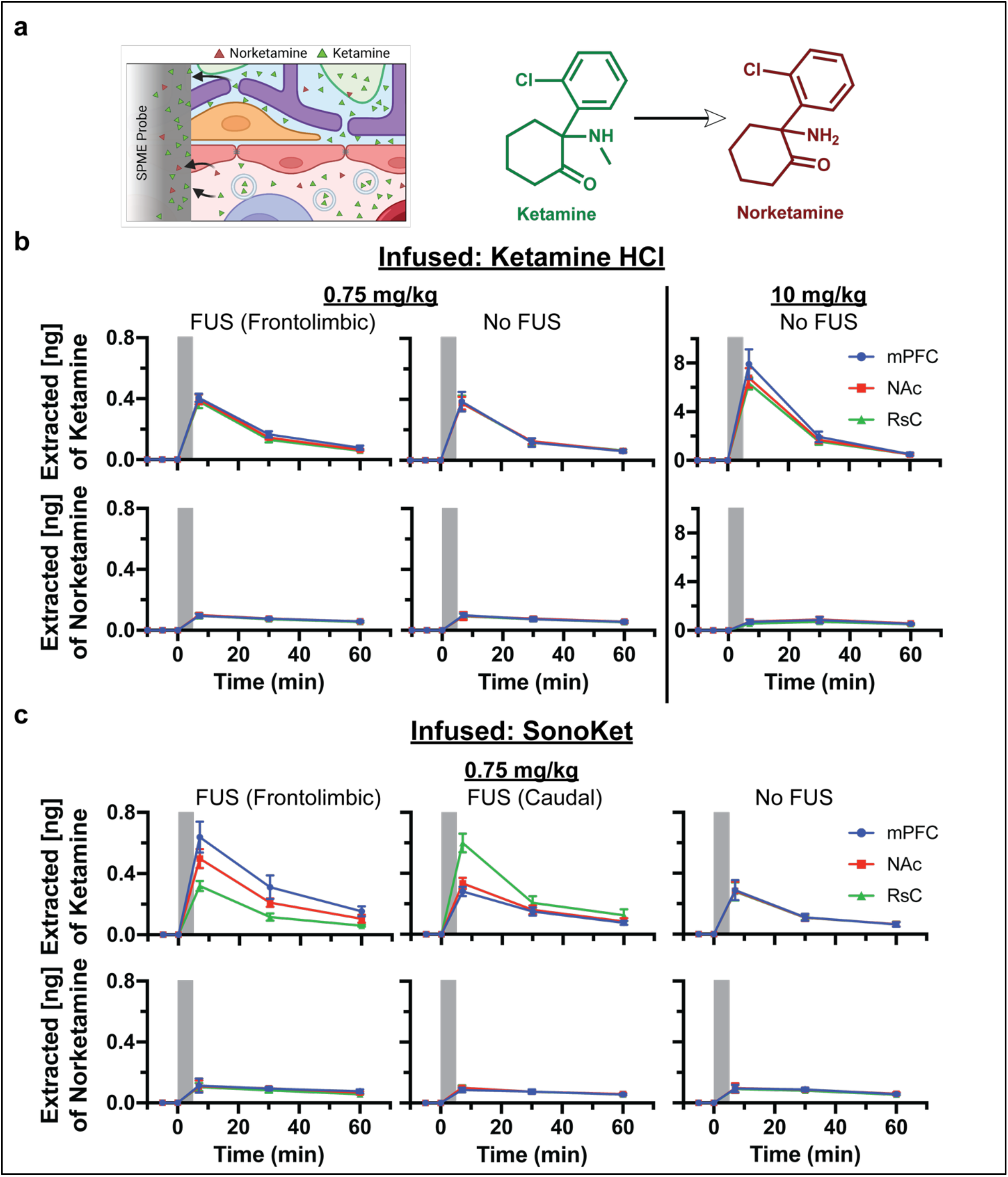
Pharmacodistribution of ketamine and norketamine across brain regions following IV administration of ketamine HCl or SonoKet, with or without FUS. (a) Molecular structures of ketamine and its metabolite norketamine (right), and a visual representation of SPME probe and the molecular diffusion of ketamine and its metabolite norketamine to the SPME probes, with norketamine deriving mainly from the blood and ketamine deriving from both the brain and blood. (b) Time-resolved pharmacodistribution of ketamine (top) and norketamine (bottom) in the mPFC (blue), NAc (red), and RsC (green) region, following free ketamine HCl IV infusion: 0.75 mg kg^-1^ free ketamine HCl with sonication at the frontolimbic region (left), 0.75 mg kg^-1^ free ketamine HCl without sonication (middle), and 10 mg kg^-1^ free ketamine HCl without sonication (right) (n=4 /group). Gray bar indicates the 5-min treatment window (drug administration with FUS or without FUS). (c) Time-resolved pharmacodistribution of ketamine (top) and its metabolite norketamine (bottom) in mPFC (blue), NAc (red), and RsC (green), following IV infusion of 0.75 mg kg^-1^ SonoKet with sonication at the frontolimbic region (left), sonication at the caudal region (middle), and without sonication (right) (*n* = 4 per group). The grey bar indicates a 5 min duration of treatment (drug administration with FUS or without FUS). Error bars represent as mean and error ± standard deviation (S.D.) for each cohort (n=4).

In contrast to free ketamine, SonoKet uncaging resulted in differential delivery of ketamine to specific brain regions. When ultrasound was targeted to the frontolimbic region during 0.75 mg kg⁻¹ SonoKet infusion, ketamine levels increased selectively and substantially at the sonicated mPFC site compared to SonoKet without FUS, with a lower rise in the NAc (where the FUS intensity was less than the mPFC) and with no rise in the non-targeted RsC (Fig. 2c, top row). Conversely, when ultrasound was directed to the caudal region containing the RsC during 0.75 mg kg⁻¹ SonoKet infusion, the highest ketamine concentration was observed at the sonicated RsC site, while the non-sonicated mPFC and NAc showed no difference compared to SonoKet without FUS, indicating region-specific drug release with minimal off-target exposure (Fig. 2c, top row). Hydroxynorketamine was undetectable (Supplementary Fig. 3, bottom row), and norketamine levels remained low across SonoKet conditions and showed no regional variation (Fig. 2c, bottom row). Notably, the peak and total levels of ketamine delivery with SonoKet uncaging were similar to that achieved with dose-matched ketamine-HCl. These results indicate that ultrasound affects the pharmacodistribution of SonoKet but not free ketamine. In particular, ultrasound enhances the release of ketamine from SonoKet selectively at the sonicated site, consistent with our prior results [25].

### Ultrasound potentiates ketamine-induced neurotransmitter and neuromodulator responses

#### Glutamate

We next examined glutamate dynamics across the mPFC, NAc, and RsC (Fig 3). Saline combined with FUS produced no significant change in glutamate across all regions, indicating that FUS alone did not induce a substantial glutamatergic modulation under these experimental conditions (Fig. 3b, top row). In contrast, free ketamine increased glutamate in a dose-dependent manner. At 0.75 mg kg⁻¹, free ketamine HCl without FUS produced a modest glutamate increase, particularly in the mPFC and NAc. When the same dose was combined with FUS targeted to the frontolimbic region, glutamate levels increased significantly more than with free ketamine alone, most prominently in the mPFC and NAc (Fig. 3b, middle row), indicating a spatially specific modulation of ketamine-induced glutamate transmission with FUS. The 10 mg kg⁻¹ free ketamine HCl (no FUS) cohort produced the largest systemic glutamate response, with prominent increases in the mPFC and NAc and a smaller response in the RsC (Fig. 3b, middle row).

**Figure 3:**
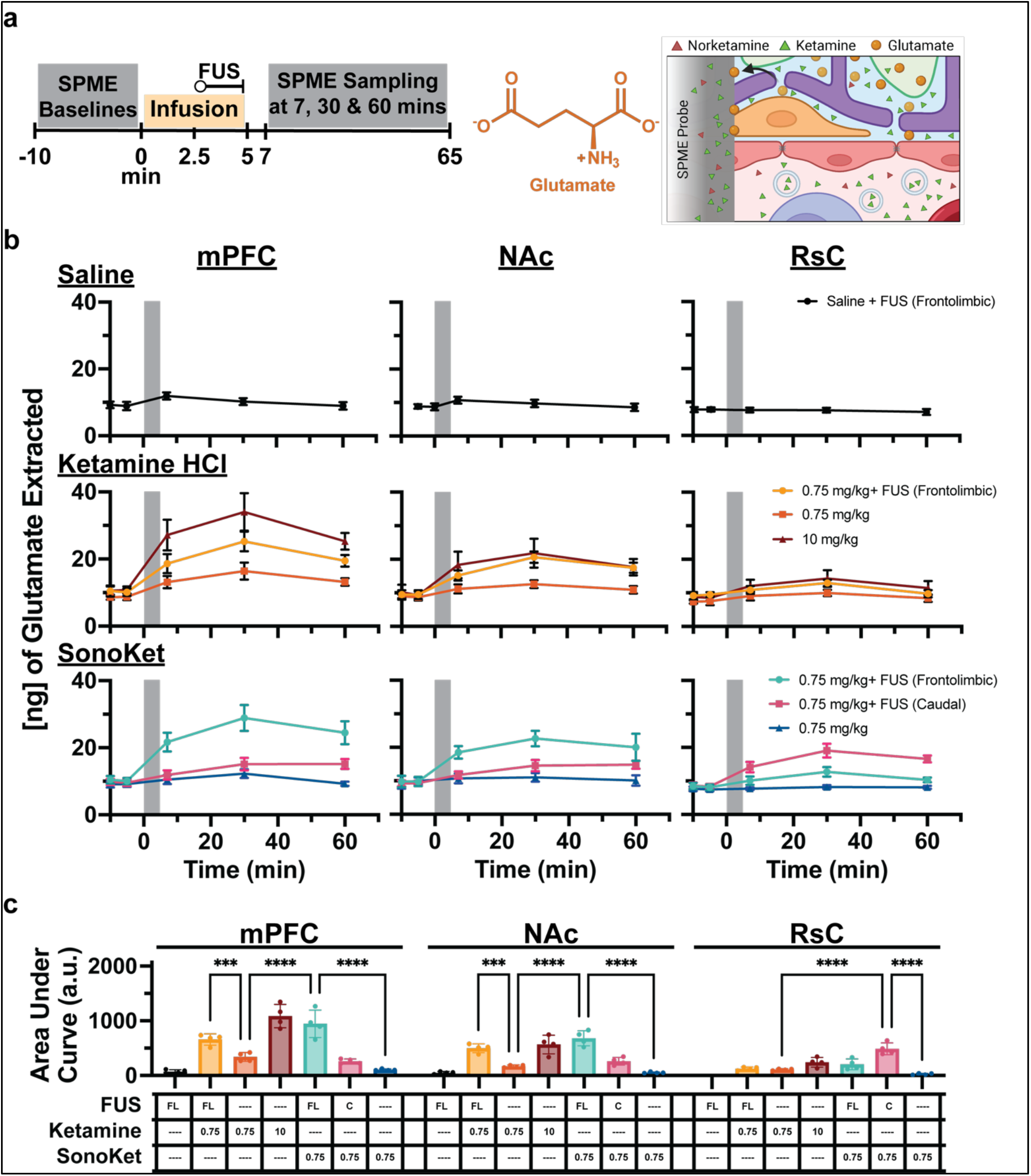
Time course dynamics of glutamate across brain regions under different ketamine and ultrasound treatment conditions. (a) Experimental timeline indicating baseline SPME sampling, treatment window and post-treatment SPME sampling (left) and molecular structure of glutamate (middle), and a schematic illustration of SPME probe and the molecular diffusion of glutamate from brain tissue and extracellular space to the SPME probe, emphasizing its neuronal origin with no direct blood contribution. (b) Time-resolved quantification of glutamate across the mPFC (left), NAc (middle), and RsC (right) regions under the different treatment conditions. Top row: Saline control with FUS targeted to the frontolimbic region. Middle row: 0.75 mg kg^-1^ free ketamine HCl with FUS targeted to the frontolimbic region, 0.75 mg kg^-1^ free ketamine HCl without FUS, and 10 mg kg^-1^ free ketamine HCL without FUS. Bottom row: 0.75 mg kg^-1^ SonoKet with FUS targeted to the frontolimbic region, 0.75 mg kg^-1^ SonoKet with FUS targeted to the caudal region, and 0.75 mg kg^-1^ SonoKet without FUS. The grey bar indicates a 5 min duration of treatment (drug administration with or without FUS). Error bars represent the mean and error ± S.D. for each cohort (*n =* 4). (c) Area under the curve (AUC) analysis quantifies the total glutamate extracted over time in each treatment group and brain region, with table indicating the experimental conditions for each bar. The FUS-treated cohorts significantly enhance glutamate levels in a region-specific manner compared to their control cohorts. ANOVA with Tukey’s multiple comparisons test. Mean ± S.D. (***p < 0.001, ****p < 0.0001).

SonoKet uncaging produced a distinct spatial and temporal glutamate profile. SonoKet without FUS produced no significant changes in glutamate compared to baseline, indicating that the potential ketamine leak from SonoKet did not impact glutamate dynamics. In contrast, SonoKet with FUS targeting the frontolimbic region resulted in a significantly greater increase in mPFC glutamate that persisted across subsequent sampling windows. Glutamate levels in the mPFC were significantly higher in the SonoKet + frontolimbic FUS condition compared to either 0.75 mg kg⁻¹ free ketamine HCl condition, with or without frontolimbic FUS. In other words, FUS potentiates the effects on glutamate concentrations of ketamine alone, with synergistically greater effects after spatially targeted uncaging. A smaller increase of glutamate was observed in the NAc after frontolimbic uncaging of SonoKet, while the RsC showed no significant change (Fig. 3b, bottom row). Conversely, when FUS was targeted to the caudal region (that includes RsC) during SonoKet infusion, the largest glutamate increase was observed in the RsC, with no significant changes in the mPFC or NAc (Fig. 3b, bottom row). Furthermore, glutamate concentrations in the RsC were significantly higher after caudal SonoKet uncaging compared to frontolimbic SonoKet uncaging and to 0.75 mg kg⁻¹ free ketamine HCl with or without frontolimbic FUS. This finding provides further evidence that SonoKet induces local spatially-specific potentiation of ketamine-induced glutamate release in the sonicated region. Overall, FUS enhanced the glutamatergic response to free ketamine, and FUS-triggered SonoKet uncaging further amplified glutamate release at the targeted region.

#### GABA

We then measured GABA to determine whether the glutamatergic response was accompanied by changes in inhibitory neurotransmission (Fig. 4). Saline combined with FUS did not affect GABA levels across the mPFC, NAc, or RsC (Fig. 4b, top row). Free ketamine HCl at either 0.75 mg kg⁻¹ or 10 mg kg⁻¹ did not produce a significant change in GABA levels; instead, levels remained near baseline or showed a non-significant trend towards a transient decrease during the early post-treatment period, most notably after 10 mg kg⁻¹ ketamine HCl, before returning toward baseline at later time points (Fig. 4b, middle row).

**Figure 4:**
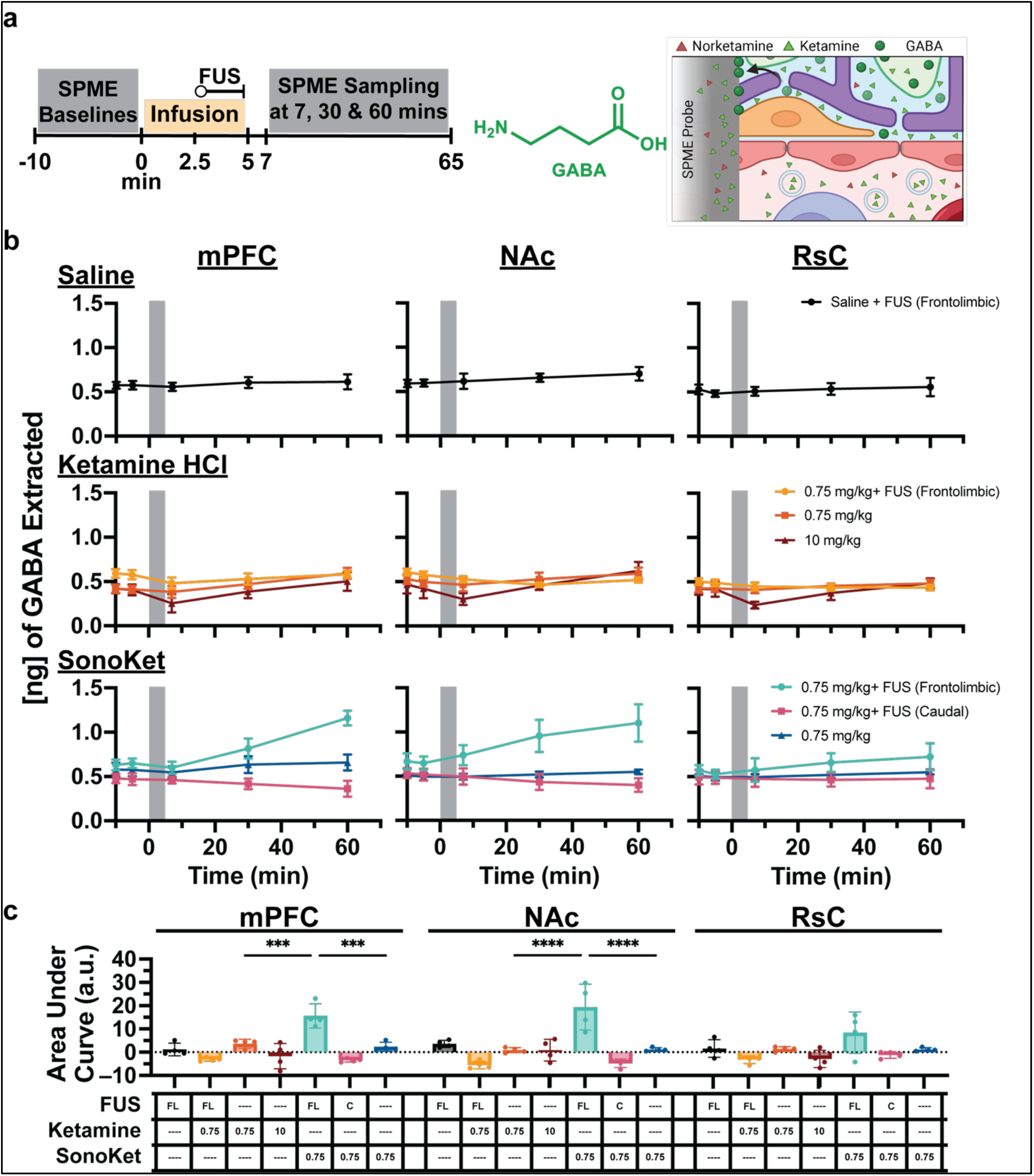
Time-resolved GABA responses across brain regions under different ketamine and ultrasound treatment conditions. (a) Experimental timeline indicating baseline SPME sampling, treatment window and post-treatment SPME sampling (left); molecular structure of γ-aminobutyric acid (GABA) (middle); and schematic illustration of SPME sampling showing the molecular diffusion of GABA from brain tissue and extracellular space along with the diffusion of ketamine and norketamine, emphasizing the origin of GABA as a neurotransmitter with no direct blood contribution (right). (b) Time-resolved dynamics of GABA across the mPFC, NAc, and RsC regions under the different treatment conditions. Top row: saline control with FUS targeted to the frontolimbic region. Middle row: 0.75 mg kg^-1^ free ketamine HCl with FUS targeted to the frontolimbic region, 0.75 mg kg^-1^ free ketamine HCl without FUS, and 10 mg kg^-1^ free ketamine HCl without FUS. Bottom row: 0.75 mg kg^-1^ SonoKet with FUS targeted to the frontolimbic region, 0.75 mg kg^-1^ SonoKet with FUS targeted to the caudal region, and 0.75 mg kg^-1^ SonoKet without FUS. The grey bar indicates a 5 min duration of treatment (drug administration with or without FUS). Error bars represent mean ± S.D. for each cohort (n=4). (c) Area under the curve (AUC) analysis quantifies the total GABA extracted over time in each treatment group and brain region, with table indicating the experimental conditions for each bar. Frontolimbic SonoKet uncaging significantly enhances GABA levels in the mPFC and NAc compared to other cohorts. ANOVA with Tukey’s multiple comparisons test. Mean ± S.D (***p < 0.001, ****p < 0.0001).

In contrast, SonoKet produced a distinct GABA profile that depended on the FUS target. SonoKet administered without FUS did not substantially alter GABA across any region indicating no pharmacologically-relevant ketamine leak with SonoKet infusion alone with respect to GABA dynamics. However, when SonoKet was uncaged with frontolimbic FUS, a delayed increase in GABA was observed at the 30–35 and 60–65 min sampling windows, most pronounced in the sonicated mPFC and NAc (Fig. 4b, bottom row). GABA levels in both the mPFC and NAc after frontolimbic SonoKet uncaging were significantly higher than GABA levels after saline with FUS, either dose of free ketamine HCl without FUS, free ketamine HCl with frontolimbic FUS, SonoKet alone, and SonoKet uncaging in the caudal region. In contrast, caudal SonoKet uncaging did not produce a comparable GABA increase in the RsC (Fig 4b, bottom row), but instead mPFC and NAc GABA levels were trend-wise lower after caudal SonoKet uncaging compared to frontolimbic SonoKet uncaging. Modulation of GABA after SonoKet uncaging is therefore region-specific. Overall, neither free ketamine nor free ketamine with FUS altered GABA; meanwhile SonoKet uncaging increased GABA release in the mPFC and NAc when FUS was directed to frontolimbic circuits and trend-wise decreased GABA release in the mPFC and NAc when FUS was directed to caudal circuits.

#### Serotonin

We next quantified serotonin (5-HT) across the same brain regions under the same conditions (Fig. 5). Saline combined with FUS produced no significant changes in 5-HT levels compared to baseline across all regions (Fig. 5b, top row). Free ketamine HCl at 0.75 mg kg⁻¹ produced a modest 5-HT increase in the mPFC and NAc, but this was not sustained; 5-HT levels returned to baseline or lower within 30 minutes after infusion. Free ketamine HCl with frontolimbic FUS not only enhanced but also sustained increases in 5-HT in the mPFC and NAc, especially the former region; 5-HT levels in the mPFC were significantly higher after 0.75 mg kg⁻¹ free ketamine HCl with frontolimbic FUS compared to 5-HT levels after 0.75 mg kg⁻¹ free ketamine HCl without FUS and saline with FUS (Fig. 5b, middle row). Intriguingly, the 10 mg kg⁻¹ free ketamine HCl group (no FUS) displayed a delayed 5-HT increase, most evident at the 30–35 min sampling mark, which was similar in peak magnitude to the increase in 5-HT after 0.75 mg kg⁻¹ free ketamine HCl with frontolimbic FUS. The RsC exhibited comparatively weaker 5-HT responses across all treatment groups.

**Figure 5.**
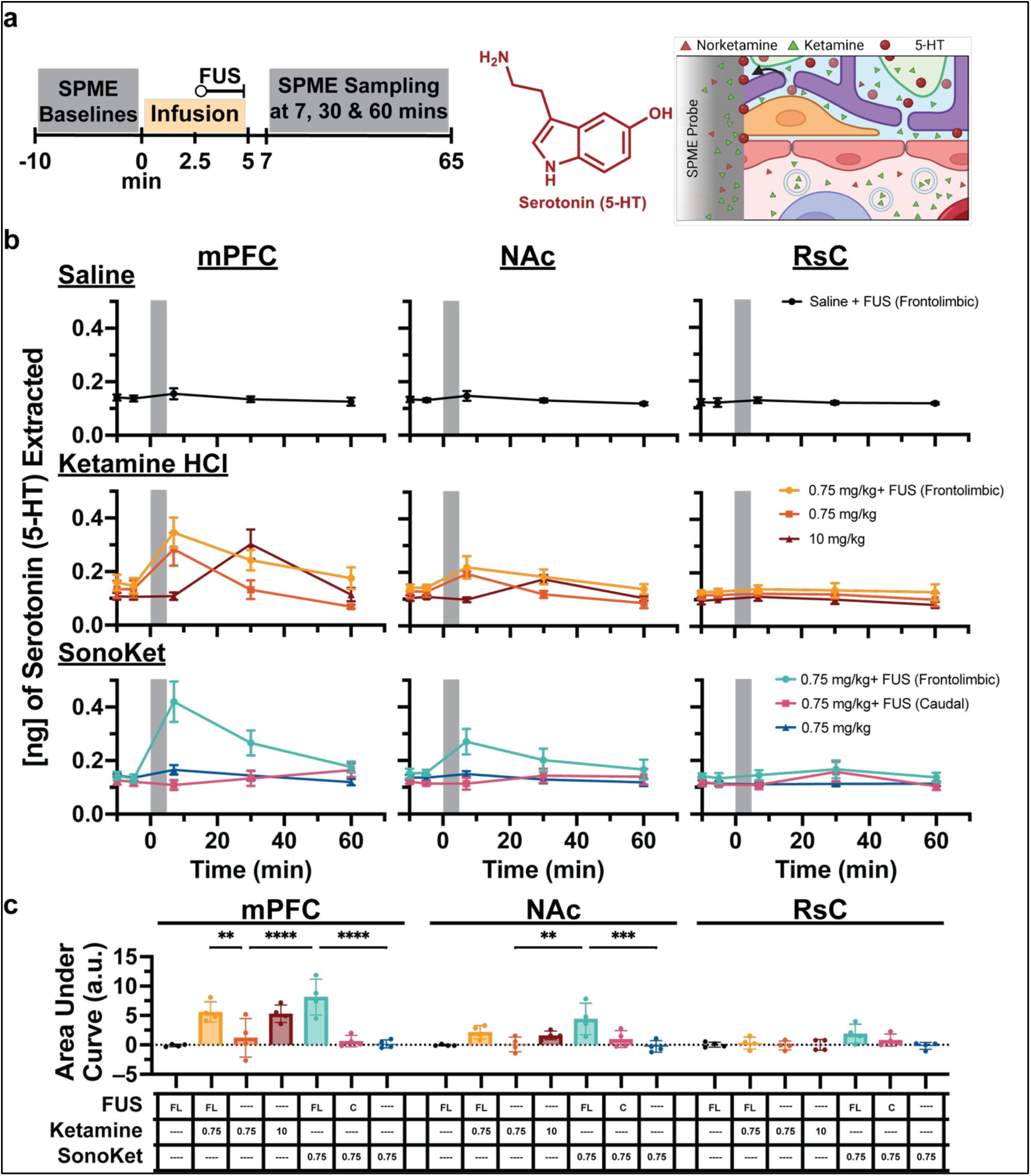
Serotonin dynamics across brain regions under different ketamine and ultrasound treatment conditions. (a) Experimental timeline indicating the sequence of SPME sampling and treatment (left); molecular structure of serotonin (middle); and schematic showing passive diffusion of 5-HT from brain tissue and extracellular space into the SPME probes along with the diffusion of ketamine and norketamine, illustrating the origin of 5-HT as a neurotransmitter with no direct blood contribution (right). (b) Time-course profiles of extracted 5-HT across the mPFC, NAc, and RsC regions under the different treatment conditions. Top row: saline control with FUS applied to the frontolimbic region. Middle row: 0.75 mg kg^-1^ free ketamine HCl with FUS targeting the frontolimbic region, 0.75 mg kg^-1^ free ketamine HCl without FUS, and 10 mg kg^-1^ free ketamine HCl without FUS. Bottom row: 0.75 mg kg^-1^ SonoKet with FUS targeting the frontolimbic region, 0.75 mg kg^-1^ SonoKet with FUS targeting the caudal region, and 0.75 mg kg^-1^ SonoKet without FUS. The grey bar indicates a 5 min duration of treatment (drug administration with or without FUS). Error bars represent mean ± S.D. for each cohort (*n =* 4). (c) Area under the curve (AUC) analysis quantifies cumulative 5-HT levels over the full post-treatment sampling period, with table indicating the experimental conditions for each bar. Combining FUS and ketamine significantly increased 5-HT in the mPFC relative to free ketamine alone, with further increased with uncaging SonoKet with frontolimbic FUS. ANOVA with Tukey’s multiple comparisons test. Mean ± S.D (**p < 0.01, ****p < 0.0001).

SonoKet frontolimbic uncaging was associated with a sharp rise in 5-HT in both the mPFC and NAc during the initial post-treatment window, while the RsC levels remained unchanged (Fig. 5b, bottom row). 5-HT levels in the mPFC after SonoKet frontolimbic uncaging were significantly higher than 5-HT levels in the mPFC after 0.75 mg kg⁻¹ free ketamine HCl without FUS, SonoKet alone, SonoKet caudal uncaging, and saline with FUS. With caudal SonoKet uncaging, a much smaller and more delayed 5-HT response was observed across sampling regions, with a modest insignificant increase at the RsC at 30–35 min. The lack of significant change in 5-HT levels with SonoKet infusion alone (without FUS) confirms the lack of pharmacologically-relevant leak of ketamine from SonoKet with respect to 5-HT dynamics. In summary, these findings indicate that free ketamine transiently elevates 5-HT in the mPFC, that frontolimbic ultrasound potentiates this increase, and that SonoKet frontolimbic uncaging leads to synergistically further potentiated levels of 5-HT in the mPFC.

#### Dopamine

Among the measured neurotransmitters, dopamine exhibited the most regionally restricted and spatially selective response (Fig. 6). Saline combined with FUS did not produce detectable dopamine levels in the mPFC, NAc, or RsC at any sampling time point, including baseline (Fig. 6b, top row). Free ketamine HCl, with or without FUS, did not yield detectable dopamine in the mPFC or RsC, but produced measurable dopamine increases in the NAc. Surprisingly, 0.75 mg kg⁻¹ free ketamine HCl (no FUS) elicited significantly more dopamine release than 10 mg kg⁻¹ free ketamine HCl (Fig. 6b, middle row). Dopamine levels in the NAc after 0.75 mg kg⁻¹ free ketamine HCl with frontolimbic FUS were slightly yet significantly higher than dopamine levels in the NAc after the same dose of free ketamine HCl without FUS.

**Figure 6:**
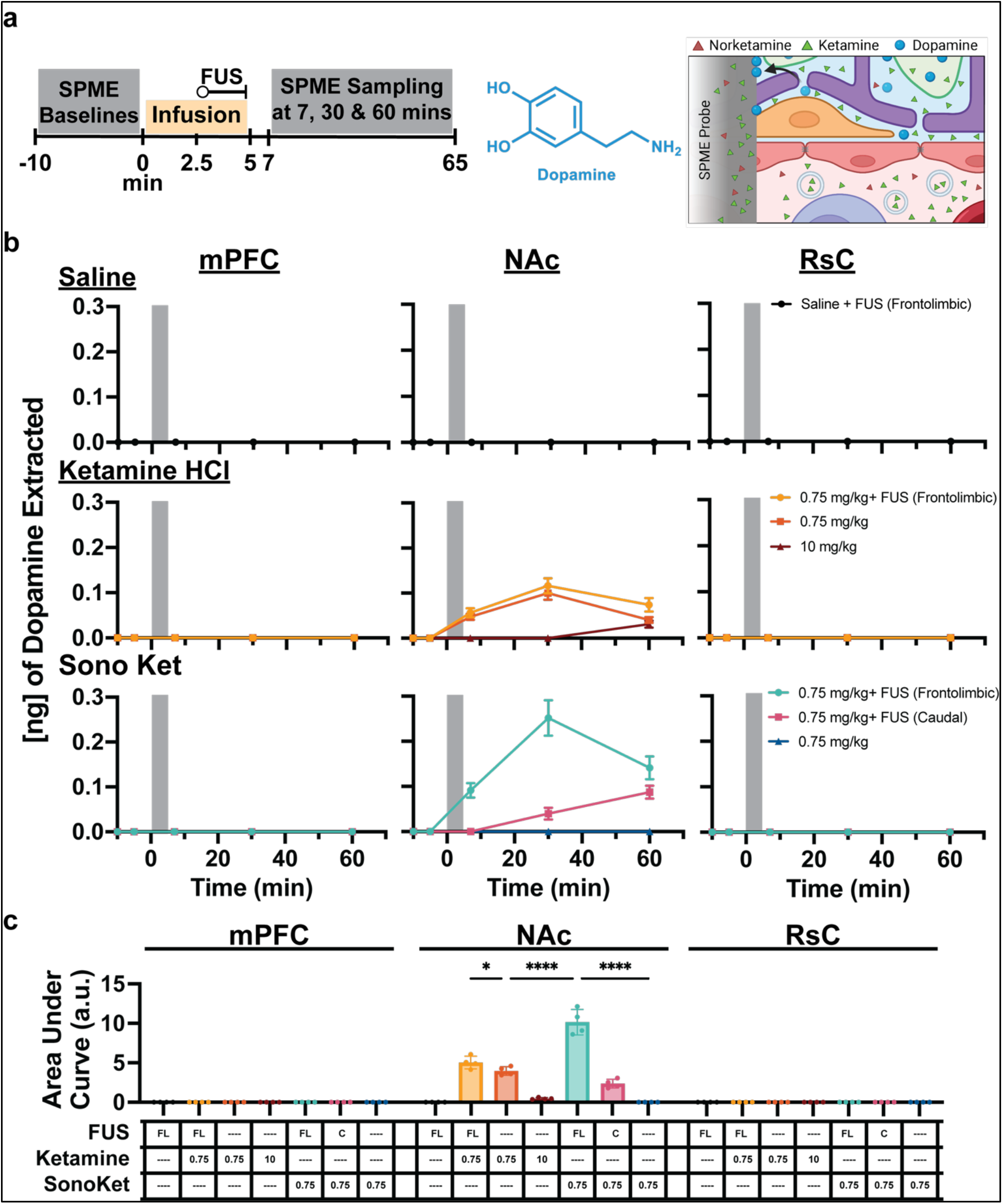
Region-specific dopamine response across the brain under different ketamine and ultrasound treatment conditions. (a) Experimental timeline with the sequence of SPME sampling and treatment (left); molecular structure of dopamine (middle); schematic showing passive diffusion of dopamine from brain tissue and extracellular space into the SPME probe along with the diffusion of ketamine and the metabolite norketamine, illustrating the origin of dopamine as a neurotransmitter with no direct blood contribution (right). (b) Time-course profiles of extracted dopamine measured in the mPFC, NAc, and RsC regions across treatment groups. Top row: saline controls with FUS targeted to the frontolimbic region. Middle row: 0.75 mg kg^-1^ free ketamine HCl with FUS targeting the frontolimbic region, 0.75 mg kg^-1^ free ketamine HCl without FUS, and 10 mg kg^-1^ free ketamine HCl without FUS. Bottom row: 0.75 mg kg^-1^ SonoKet with FUS targeted to the frontolimbic region, 0.75 mg kg^-1^ SonoKet with FUS targeting the caudal region, and 0.75 mg kg^-1^ SonoKet without FUS. Notably, dopamine levels remained largely undetectable in the mPFC and RsC regions, irrespective of any treatment group. <u>The grey bar indicates a 5 min duration of treatment (drug administration with or without FUS).</u> Error bars represent mean ± S.D. for each cohort (*n* = 4). (c) Area under the curve (AUC) analysis quantifies collective dopamine responses over the full post-treatment sampling period, with table indicating the experimental conditions for each bar. Dopamine is significantly elevated in NAc with frontolimbic SonoKet uncaging, compared to all other conditions. ANOVA with Tukey’s multiple comparisons test. Mean ± S.D (*p < 0.05, ****p < 0.0001).

SonoKet infusion without FUS produced no measurable changes in dopamine levels, confirming that leak of ketamine from SonoKet does not significantly affect dopamine dynamics. SonoKet uncaging at either the frontolimbic or caudal regions produced a dopamine response exclusively in the NAc, with no detectable levels of dopamine in the mPFC or RsC across any SonoKet treatment group. With frontolimbic SonoKet uncaging, NAc dopamine increased markedly, peaking during the 30–35 min sampling window and declining by 60–65 min, yet remaining elevated relative to baseline and the initial post-treatment time point (Fig. 6b, bottom row). Increases in dopamine were significantly higher after SonoKet frontolimbic uncaging relative to all other conditions. In contrast, caudally targeted SonoKet uncaging produced a smaller, delayed NAc dopamine increase. These results indicate that low doses of free ketamine raise dopamine levels in the NAc, and both FUS and frontolimbic-targeted SonoKet uncaging further potentiates dopamine release. This potentiation occurs selectively in the NAc, rather than globally across sampled regions.

Of note, these results may underestimate dopamine levels in each region and could be relatively insensitive to more modest changes in dopamine levels due to potential oxidation of the dopamine species during extraction and sample preparation that may reduce sensitivity to this neurotransmitter.

### Restraint acclimatization preserves FUS-induced ketamine neurochemical responses

Awake restraint can induce an acute stress state in rodents [40,41], which may influence neurochemical profiles during *in vivo* sampling. To determine whether acute stress from awake restraint affected the neurochemical response to ketamine with or without ultrasound, rats were acclimatized in the restraint apparatus for 90 min per day on three consecutive days before the experiment. During the experiment, rats received IV infusion of low-dose free ketamine HCl (0.75 mg kg⁻¹) for 5 min, with ultrasound applied during the final 2.5 min, followed by SPME sampling in the mPFC, NAc, and RsC.

As shown in Supplementary Fig. 4, ketamine levels increased rapidly in all sampling regions and declined by 60–65 min, while norketamine remained lower throughout, with no significant differences compared to the results acquired without restraint acclimatization (Fig. 2). Despite the widespread distribution of ketamine across the brain, neurotransmitter responses showed regional selectivity. Glutamate increased most prominently in the mPFC and NAc, while the RsC response remained comparatively stable. 5-HT also showed an early increase in the mPFC and NAc, whereas GABA levels remained relatively stable across all regions and time points. Dopamine showed the most restricted response, increasing selectively in the NAc and peaking at 30–35 min, with no detectable levels in the mPFC or RsC. These findings reproduce the regional neurochemical patterns observed in the main experimental cohorts without restraint acclimatization (Fig. 2–6), indicating that acute stress from restraint did not substantially alter these neurochemical levels and that FUS-potentiated ketamine responses were preserved under acclimatization conditions.

## Discussion

In this study we demonstrated that FUS and ketamine interact to alter brain neurochemistry in a spatially specific manner – with synergistically greater effects with localized ketamine delivery via ultrasonic uncaging. Using *in vivo* SPME sampling followed by LC-MS/MS analysis, we showed that FUS alone does not detectably alter glutamate, GABA, serotonin, or dopamine levels across the mPFC, NAc, or RsC, nor does it significantly alter the regional pharmacodistribution of free ketamine or its metabolites. Although prior work reported changes in extracellular GABA, dopamine, and serotonin after applying FUS to the rodent brain, these studies used much higher pulse repetition frequencies as well as a different sampling methodology, microdialysis, which may have differential sensitivity to regionally restricted release of neurotransmitters [42–44]. To our knowledge, this is the first study to apply SPME to investigate FUS neuromodulation *in vivo*, with each SPME fiber capturing ketamine, its metabolites, glutamate, GABA, 5-HT, and dopamine in a single extraction. This approach enabled multiplexed measurement of regional drug exposure and neurotransmitter dynamics from the same brain site, allowing FUS-targeted SonoKet uncaging to be distinguished from non-specific leakage or systemic free ketamine exposure. By placing fibers concurrently in the mPFC, NAc, and RsC of the same animal, we were able to directly relate region-specific ketamine pharmacokinetics and pharmacodistribution to downstream neurochemical responses across all three regions in parallel.

While FUS alone did not change the concentrations of neurotransmitters and neuromodulators, pairing FUS with ketamine significantly augmented glutamate, serotonin, and dopamine compared to free ketamine administration alone. The fact that FUS only enhanced neurotransmitter concentrations in the presence of ketamine indicates that FUS does not directly interact with cellular mechanisms that mediate the release of these neurotransmitters, such as vesicular release or monoamine transporters. Instead, FUS’ neuromodulatory effects appear to be dependent on the underlying state of the sonicated circuit, similar to demonstrated results of FUS modulation of TMS- or DBS-induced activities [45,46]. FUS only increases neurotransmitter release at a circuit when that circuit has already been engaged with a stimulus like ketamine or electrical modulation. It is possible that ketamine pushes neural circuits into high-gain regimes in which FUS’ typically subthreshold effects become detectable. As a potential mechanism for these findings, Yoo et al. demonstrated that focused ultrasound activates mechanosensitive calcium-permeable channels, producing a gradual accumulation of calcium. Oh et al. showed that ultrasound also induces mechanosensitive channel activation in astrocytes, which increases the influx of calcium in neurons. Both of these processes can be amplified by voltage-gated sodium and calcium channels to generate burst firing [47,48]. However, under baseline conditions, the excitation-inhibition ratio may be too low for calcium accumulation to reach the threshold for voltage-gated amplification [49,50]. Subanesthetic ketamine preferentially blocks NMDA receptors on GABA interneurons, disinhibiting downstream pyramidal cells and shifting them toward a depolarized, high-gain state in which voltage-gated channels may be more readily recruited [51,52]. In turn, this may increase the probability of presynaptic release of neurotransmitters [53]. Accordingly, FUS alone may not result in enough calcium accumulation to trigger or alter action potentials, but ketamine, by depolarizing pyramidal neurons, may enable the same calcium signal to recruit voltage-gated channels and yield presynaptic transmitter release. Hence, FUS and ketamine can synergistically interact to drive up the concentrations of glutamate and other neuromodulators. In addition, FUS may alter the phenotype of glial cells [48,54] that normally serve to maintain the synaptic cleft, which may contribute to the observed effects. This suggests that the therapeutic window for FUS-based interventions may be substantially widened by co-administration with a pharmacological agent that primes the targeted circuit.

Beyond the co-administration of ketamine and FUS, uncaging ketamine via SonoKet further enhances the concentration of glutamate, GABA, serotonin, and dopamine at the sonicated region. In other words, targeting ketamine to specific brain regions with ultrasonic uncaging is even more potent than ultrasound and free ketamine, or dose-matched free ketamine alone, at driving neurotransmitter and neuromodulator release. In fact, glutamate levels in the mPFC after frontolimbic 0.75 mg kg^-1^ SonoKet uncaging were roughly equivalent to those elicited by 10 mg kg^-1^ free ketamine. The enhanced neuromodulator release after SonoKet uncaging may explain the potent increases in gamma power and changes in rodent behavior that were greater than dose-matched free ketamine observed in our prior studies [25]. Notably, the lack of change of any neurotransmitter level with SonoKet infusion alone confirms that there is no pharmacologically relevant leak of ketamine from SonoKet *in vivo* under these conditions.

While SonoKet uncaging elevated all four of the assessed neurotransmitters and neuromodulators, some effects were unique to particular chemicals. Glutamate responses were the most spatially constrained, with the largest increases co-localized to the FUS-targeted region (mPFC or NAc with frontolimbic uncaging, RsC with caudal uncaging). This pattern is consistent with direct local engagement of pyramidal neurons at the FUS focus and propagation through known mPFC efferents to the NAc [55]. GABA followed a similar regional logic but with a more delayed and smaller magnitude of change, in line with its role as a feedback signal recruited downstream of pyramidal-cell activation rather than as a primary driver of the response [56,57]. Serotonin showed the most temporally extended elevation, with frontolimbic SonoKet uncaging producing 5-HT increases in the mPFC and NAc that were sustained well beyond the early post-treatment window, in contrast to the transient 5-HT response observed with dose-matched free ketamine alone. This sustained 5-HT signal may reflect engagement of slower neuromodulatory loops, possibly via mPFC projections to the dorsal raphe and reciprocal serotonergic innervation of frontal and striatal targets, which integrate over longer timescales than the fast glutamatergic and GABAergic responses [58–60]. Finally, dopamine showed the most regionally restricted pattern of all four analytes. Despite ketamine being widely distributed across all three sampled regions, dopamine was detectable only in the NAc, likely because the density of dopaminergic terminals in mPFC and RsC is substantially lower than that of the NAc, and due to a potential technical limitation of dopamine oxidation reducing sensitivity to low dopamine levels [61–63].

Overall, with uncaging the ketamine levels achieved at the target site were only marginally greater than those achieved with dose-matched free ketamine. Compared to dose-matched free ketamine the most prominent difference is that the rest of the brain and body outside the sonicated region is exposed to less ketamine. That uncaging yields even greater functional effects than ultrasound with free ketamine suggests that center-surround effects seen with visual and electrical stimuli also apply to pharmacologic stimuli.

Together, these findings establish that focused ultrasound and ketamine interact in a synergistic and spatially selective manner to reshape neurochemical dynamics across functionally distinct brain circuits. FUS alone does not detectably alter extracellular neurotransmitter concentrations with the ultrasound parameters used here, but it substantially amplifies the neurochemical response to ketamine when the two are administered together. SonoKet uncaging, which deposits ketamine at the ultrasound focus, produces the largest neuromodulatory response, even at a fraction of the systemic dose required for comparable effects with free ketamine. Translationally, the regional specificity of SonoKet uncaging combined with the order-of-magnitude reduction in systemic ketamine exposure suggests that SonoKet can potentially optimize the therapeutic actions of ketamine while minimizing its systemic toxicities and dissociative and abuse-related liabilities. Overall, these results support that ultrasonic drug uncaging offers a generalizable framework for circuit-targeted pharmacotherapy.

## Methods

### Materials

All chemicals and reagents used in this study were of the highest purity grade. Ketamine-HCl injectable solution (100 mg mL-1), a controlled substance, was obtained through Stanford University Environmental Health & Safety. Liposomal ketamine, termed ‘SonoKet’, was synthesized in-house using a method previously reported by our group [25]. HPLC-and LC/MS-grade water, methanol, 2-propanol, acetonitrile, and formic acid were purchased from Fisher Scientific. Cerilliant® certified reference solutions of ketamine hydrochloride, norketamine hydrochloride, hydroxynorketamine hydrochloride, serotonin hydrochloride (5-HT), and dopamine hydrochloride (DA), along with the deuterated internal standards ketamine-D4 hydrochloride, norketamine-D4 hydrochloride, serotonin-D4 hydrochloride, and dopamine-D4 hydrochloride, were obtained from MilliporeSigma (USA). Standards of γ-aminobutyric acid (GABA) and glutamic acid (Glu) were also obtained from MilliporeSigma, whereas their isotopically labelled analogues (GABA-D6 and glutamic acid-D5, used as internal standards) were sourced from LGC Standards (USA). Custom C8-SCX/PAN SPME fibers were procured from MilliporeSigma (Bellefonte, PA, USA). Biocompatible sterile Kwik-Sil/Kwik-Cast silicone surgical bioadhesive was obtained from World Precision Instruments (FL, USA).

### Animals

All animal experiments were conducted in accordance with the Stanford Institutional Animal Care and Use Committee (IACUC) and Administrative Panel on Laboratory Animal Care (APLAC). Male Sprague-Dawley rats aged 8–10 weeks with body weights of approximately 300– 400 g (Charles River Laboratories, Wilmington, MA; Envigo, Indianapolis, IN) were used in all *in vivo* studies involving ketamine HCl and SonoKet. Isoflurane anaesthesia was used for all surgical and terminal procedures and was administered briefly as needed during awake animal experimental preparation.

### Dose preparation

Ketamine hydrochloride (Dechra Veterinary Products) was diluted in 0.9% sterile saline to yield working solutions of 1 mg mL⁻¹ and 10 mg mL⁻¹. In-house synthesized SonoKet was diluted in 0.9% sterile saline to 1 mg mL⁻¹. All treatments were administered via intravenous tail-vein infusion over 5 min using a syringe infusion pump (World Precision Instruments).

### Animal preparation

One day before treatment, rats were anaesthetized, placed in a stereotaxic frame, and maintained on a heating pad at 37°C. Each animal received 2 mL saline and buprenorphine (0.05 mg kg⁻¹) subcutaneously for preoperative care. A midline scalp incision was made and three burr holes (∼1 mm diameter) were drilled into the skull under stereotaxic guidance, centered at the following coordinates (relative to bregma): mPFC, +3.2 mm A/P, +0.8 mm M/L; NAc, +1.6 mm A/P, +0.8 mm M/L; RsC, −4.0 mm A/P, +0.8 mm M/L (Fig. 1) [64]. A durotomy was performed at each site using a 32 G needle to facilitate SPME probe insertion for direct sampling of drugs and neurochemicals in the brain immediately after treatment. Following surgery, scalp wounds were sealed with biocompatible sterile Kwik-Sil/Kwik-Cast silicone bioadhesive, and animals were housed individually and allowed to recover for at least 24 hours before the treatment and SPME experiment.

### *In vivo* experiment and solid-phase microextraction (SPME)

Before each experiment, SPME fibers were cleaned with methanol:acetonitrile:isopropanol (2:1:1, v/v/v), preconditioned overnight in methanol:water (1:1, v/v), rinsed with water and stored in water until use. At the start of each experiment, rats were briefly anaesthetized for tail-vein catheterization, placed in a thin flexible plastic restraint cone, and secured in a custom head-restraining apparatus as previously described [65]. The scalp bioadhesive seal was removed and animals were provided supplemental oxygen via nose cone. A 30 min equilibration period was allowed after restraint placement to ensure complete isoflurane washout and full return to the awake state before initiation of treatment and SPME sampling.

For animals receiving focused ultrasound (FUS), the transducer was positioned directly above the target burr hole over either the mPFC (+3.2 mm A/P, +0.8 mm M/L) or the RsC (−4.0 mm A/P, +0.8 mm M/L) using a three-axis positioning system (ThorLabs). Acoustic coupling was achieved with a coupling cone and ultrasound gel. Ultrasound was delivered using custom-built transducers at 250 kHz calibrated with hydrophones (Onda) under the following parameters: 50 ms ON, 150 ms OFF (25% duty cycle), 5 Hz pulse repetition frequency. Acoustic parameters were set to deliver a target in situ peak negative pressure of 1.1 MPa at the focal point, with transcranial attenuation estimated based on animal weight and center frequency following O’Reilly et al. [66].

For SPME sampling, fibers were inserted 3 mm below the brain surface through the burr holes, positioning the extraction phase centered approximately 3.0 mm below the brain surface. Fibers were exposed to tissue for 5 min to allow diffusion-based extraction of ketamine, its metabolites, and endogenous neurochemicals onto the fiber coating, consistent with SPME principles [27–32]. Two baseline samples were collected before treatment, followed by three post-treatment collections at 2–7 min, 25–30 min, and 55–60 min. Treatment groups (n = 4 per group) were: saline vehicle; ketamine HCl (0.75 mg kg⁻¹) with or without FUS; ketamine HCl (10 mg kg⁻¹) without FUS; and SonoKet (0.75 mg kg⁻¹) with or without FUS. Ultrasound was applied during the final 2.5 min of the 5 min IV infusion. Following sampling, SPME fibers were withdrawn, gently wiped to remove residual blood, and desorbed into 50 μL of MeOH/H₂O (7:3, v/v) containing 0.1% formic acid and deuterated internal standards for 30 min at room temperature under agitation at 1500 rpm. The resulting eluates were quantified by LC–MS/MS using an instrumental calibration curve.

### LC–MS/MS analysis

LC–MS/MS analysis was performed using an Agilent 1290 Infinity LC System coupled to an Agilent 6490 Triple Quadrupole LC/MS System with iFunnel technology (Agilent Technologies, San Jose, CA, USA). Analyses were conducted in positive ionization mode using Agilent Jet Stream electrospray ionization (AJS-ESI) under optimized MS/MS conditions (Supplementary Table 1). A fragmentor voltage of 380 V and a cell accelerator voltage of 5 V were applied consistently across all SRM transitions. MS source parameters are summarized in Supplementary Table 2. Data acquisition was performed using Agilent MassHunter Workstation Data Acquisition software (version B.08.00) and quantitative processing using MassHunter Workstation Quantitative Analysis software (version B.07.01; Agilent Technologies).

Chromatographic separation was performed at room temperature using a Phenomenex Kinetex PFP column (1.7 μm, 100 Å, 100 mm × 2.1 mm; part no. 00D-4476-AN) equipped with a Phenomenex SecurityGuard ULTRA UHPLC PFP cartridge (2.1 mm ID; part no. AJ0-8787). Mobile phase A consisted of LC/MS-grade water with 0.1% formic acid and 1 mM perfluoropentanoic acid; mobile phase B consisted of LC/MS-grade methanol with 0.1% formic acid and 1 mM perfluoropentanoic acid. The injection volume was 4 μL and the flow rate was 300 μL min^-1^ with a gradient as detailed in Supplementary Table 3. Column back-pressure was maintained at a maximum of 1000 bar and the autosampler temperature was set to 5°C. Analytical responses, expressed as relative peak area ratios (analyte to internal standard), were converted to extracted amounts using a calibration curve prepared in MeOH:H₂O (7:3, v/v) with 0.1% formic acid and internal standards at 50 ng mL^-1^; analyte concentrations ranged from 0.1 to 100 ng mL^-1^.

### Ultrasound simulations

Micro-CT imaging was performed on a male Sprague-Dawley rat (375 g) using a Quantum GX scanner (PerkinElmer, MA, USA). Images comprised 1,024 × 1,024 × 553 voxels at 0.086 mm isotropic resolution and were resampled to 0.34 mm for simulation. Bone, soft tissue, and water were segmented based on Hounsfield unit thresholds. Tissue density and sound speed were interpolated using the hounsfield2density function in the k-Wave MATLAB toolbox (MathWorks, Natick, MA, USA). The transducer was modelled as a 100 mm diameter bowl with a plastic coupling cone, with geometry and material properties matched to the experimental transducer setup. Simulations were run at 250 kHz with a time step of 22.8 ns for 85 μs to cover the full propagation grid.

## Statistical analysis

All data are presented as mean ± standard deviation. Within-group comparisons across multiple time points were performed using one-way repeated-measures ANOVA, with Tukey’s or Bonferroni’s post-hoc test to identify significant pairwise differences. Between-group comparisons across time were assessed using two-way repeated-measures ANOVA with group and time as factors, followed by appropriate post-hoc correction for multiple comparisons. Area under the curve (AUC) and peak response values were calculated for each animal and compared across experimental groups using one-way ANOVA with Tukey’s multiple comparisons test. Given the small sample size (n = 4 per group), normality was assessed using the Shapiro–Wilk test; where normality was violated, non-parametric alternatives were used: the Friedman test with Dunn’s post-hoc correction for repeated-measures comparisons and the Kruskal–Wallis test with Dunn’s correction for independent group comparisons. AUC was calculated using the trapezoidal method. All statistical analyses were performed in GraphPad Prism (version 10.0; GraphPad Software, San Diego, CA), and statistical significance was set at p < 0.05. See Supplementary Tables 4-6 for individual comparison p-values.

## Acknowledgments

We want to thank Mario Landeros Wences for his support and assistance with logistics and study supplies. We also thank Abel Bermudez and Dr. Sharon Petteri from the Canary Center, Stanford University, for their support in maintaining the HPLC and LC–MS/MS instruments during the study. We are grateful to Ronald Watkins for his assistance in producing the ultrasound hardware used in these experiments. Finally, we thank all previous and current members of the Airan Lab for their helpful discussions and support.

## Funding

Seed Grant from the Stanford Wu Tsai Neurosciences Institute (RDA)

NIH BRAIN Initiative (NIH/NIMH RF1MH114252, NIH/NINDS UG3NS114438 to RDA)

NIH HEAL Initiative (NIH/NINDS UG3NS115637 to RDA)

Anonymous Donor to the Stanford SOM Radiology Department (RDA)

## Authors Contributions

Conceptualization: RDA, KSR, PJM

Methodology: KSR, PJM, MMP, SNE, YX, RDA

Analysis: KSR, PJM, RDA

Visualization: KSR, PJM, RDA

Funding Acquisition: RDA

Project Administration: RDA

Supervision: RDA

Writing (Initial Draft): KSR, PJM, KS, RDA

Review and Editing: All authors

## References

[1] Legon W, Adams S, Bansal P, Patel PD, Hobbs L, Ai L, et al. A retrospective qualitative report of symptoms and safety from transcranial focused ultrasound for neuromodulation in humans. Sci Rep 2020;10:5573. 10.1038/s41598-020-62265-8.

[2] Legon W, Strohman A. Low-intensity focused ultrasound for human neuromodulation. Nature Reviews Methods Primers 2024;4:91. 10.1038/s43586-024-00368-6.

[3] Legon W, Sato TF, Opitz A, Mueller J, Barbour A, Williams A, et al. Transcranial focused ultrasound modulates the activity of primary somatosensory cortex in humans. Nat Neurosci 2014;17:322–9. 10.1038/nn.3620.

[4] Shi Y, Cai G, Wu W. A panoramic review of transcranial focused ultrasound neuromodulation: from basic research to clinical applications. J Neuroeng Rehabil 2025;22:227. 10.1186/s12984-025-01753-2.

[5] Mcmahon D, O’reilly MA, Hynynen K. Therapeutic Agent Delivery Across the Blood-Brain Barrier Using Focused Ultrasound 2026;29:54. 10.1146/annurev-bioeng-062117.

[6] Lee K, Park TY, Lee W, Kim H. A review of functional neuromodulation in humans using low-intensity transcranial focused ultrasound. Biomed Eng Lett 2024;14:407–38. 10.1007/s13534-024-00369-0.

[7] Park C, Kim SE, Choi YS, Lee J, Lee SY, Kang JH, et al. Structural-Functional Brain Network Modulation using Transcranial Focused Ultrasound Stimulation: Implications on the Default Mode Network in Humans. Neuroimage 2025;321:121540. 10.1016/j.neuroimage.2025.121540.

[8] Beisteiner R, Hallett M, Lozano AM. Ultrasound Neuromodulation as a New Brain Therapy. Advanced Science 2023;10. 10.1002/advs.202205634.

[9] Legon W, Strohman A. Low-intensity focused ultrasound for human neuromodulation. Nature Reviews Methods Primers 2024;4. 10.1038/s43586-024-00368-6.

[10] Darrow DP. Focused Ultrasound for Neuromodulation. Neurotherapeutics 2019;16:88–99. 10.1007/s13311-018-00691-3.

[11] Krystal JH, Kavalali ET, Monteggia LM. Ketamine and rapid antidepressant action: new treatments and novel synaptic signaling mechanisms. Neuropsychopharmacology 2024;49:41–50. 10.1038/s41386-023-01629-w.

[12] Krystal JH, Kaye AP, Jefferson S, Girgenti MJ, Wilkinson ST, Sanacora G, et al. Ketamine and the neurobiology of depression: Toward next-generation rapid-acting antidepressant treatments. Proc Natl Acad Sci U S A 2023;120. 10.1073/pnas.2305772120.

13. Berman RM, Cappiello A, Anand A, Oren DA, Heninger GR, Charney DS, et al. BRIEF REPORTS Antidepressant Effects of Ketamine in Depressed Patients. vol. 47. 2000.

[14] Mathew CAZSJ, Mathew SJ, Zarate Jr CA. Ketamine for treatment-resistant depression. Springer; 2016.

[15] Morgan CJA, Curran HV, (ISCD) the ISC on D. Ketamine use: a review. Addiction 2012;107:27–38. 10.1111/j.1360-0443.2011.03576.x.

[16] Sassano-Higgins S, Baron D, Juarez G, Esmaili N, Gold M. A REVIEW OF KETAMINE ABUSE AND DIVERSION. Depress Anxiety 2016;33:718–27. 10.1002/da.22536.

[17] Wolff K, Winstock AR. Ketamine From Medicine to Misuse. vol. 20. 2006.

[18] Duman RS, Aghajanian GK, Sanacora G, Krystal JH. Synaptic plasticity and depression: New insights from stress and rapid-acting antidepressants. Nat Med 2016;22:238–49. 10.1038/nm.4050.

[19] Duman RS, Li N, Liu RJ, Duric V, Aghajanian G. Signaling pathways underlying the rapid antidepressant actions of ketamine. Neuropharmacology, vol. 62, 2012, p. 35–41. 10.1016/j.neuropharm.2011.08.044.

[20] Vesuna S, Kauvar I V., Richman E, Gore F, Oskotsky T, Sava-Segal C, et al. Deep posteromedial cortical rhythm in dissociation. Nature 2020;586:87–94. 10.1038/s41586-020-2731-9.

[21] Masuzawa M, Nakao S, Miyamoto E, Yamada M, Murao K, Nishi K, et al. Pentobarbital Inhibits Ketamine-Induced Dopamine Release in the Rat Nucleus Accumbens: A Microdialysis Study. Anesth Analg 2003;96.

[22] Rizzo A, Garçon-Poca MZ, Essmann A, Souza AJ, Michaelides M, Ciruela F, et al. The dopaminergic effects of esketamine are mediated by a dual mechanism involving glutamate and opioid receptors. Mol Psychiatry 2025;30:3443–54. 10.1038/s41380-025-02931-3.

[23] Cardona-Acosta AM, Bolaños-Guzmán CA. Role of the mesolimbic dopamine pathway in the antidepressant effects of ketamine. Neuropharmacology 2023;225. 10.1016/j.neuropharm.2022.109374.

[24] Marcott PF, Gong S, Donthamsetti P, Grinnell SG, Nelson MN, Newman AH, et al. Regional Heterogeneity of D2-Receptor Signaling in the Dorsal Striatum and Nucleus Accumbens. Neuron 2018;98:575–587.e4. 10.1016/j.neuron.2018.03.038.

[25] Purohit MP, Yu BJ, Roy KS, Xiang Y, Ewbank SN, Azadian MM, et al. Acoustically activatable liposomes as a translational nanotechnology for site-targeted drug delivery and noninvasive neuromodulation. Nat Nanotechnol 2025;20:1688–99. 10.1038/s41565-025-01990-5.

[26] Pawliszyn J. Theory of Solid-Phase Microextraction. Handbook of Solid Phase Microextraction, Elsevier Inc.; 2012, p. 13–59. 10.1016/B978-0-12-416017-0.00002-4.

[27] Reyes-Garcés N, Gionfriddo E, Gómez-Ríos GA, Alam MN, Boyacl E, Bojko B, et al. Advances in Solid Phase Microextraction and Perspective on Future Directions. Anal Chem 2018;90:302–60. 10.1021/acs.analchem.7b04502.

[28] Huq M, Tascon M, Nazdrajic E, Roszkowska A, Pawliszyn J. Measurement of Free Drug Concentration from Biological Tissue by Solid-Phase Microextraction: In Silico and Experimental Study. Anal Chem 2019;91:7719–28. 10.1021/acs.analchem.9b00983.

[29] Roy KS, Nazdrajić E, Shimelis OI, Ross MJ, Chen Y, Cramer H, et al. Optimizing a High-Throughput Solid-Phase Microextraction System to Determine the Plasma Protein Binding of Drugs in Human Plasma. Anal Chem 2021;93:11061–5. 10.1021/acs.analchem.1c01986.

[30] Reyes-Garcés N, Diwan M, Boyacl E, Gómez-Ríos GA, Bojko B, Nobrega JN, et al. In vivo brain sampling using a microextraction probe reveals metabolic changes in rodents after deep brain stimulation. Anal Chem 2019;91:9875–84. 10.1021/ACS.ANALCHEM.9B01540/SUPPL_FILE/AC9B01540_SI_002.XLSX.

[31] Lendor S, Hassani SA, Boyaci E, Singh V, Womelsdorf T, Pawliszyn J. Solid Phase Microextraction-Based Miniaturized Probe and Protocol for Extraction of Neurotransmitters from Brains in Vivo. Anal Chem 2019;91:4896–905. 10.1021/acs.analchem.9b00995.

[32] Nazdrajić E, Roy KS, Jaroch K, Markuszewski MJ, Olkowicz M. The evolving role of solid-phase microextraction in translational medicine: Forging the path forward to personalized medicine. TrAC Trends in Analytical Chemistry 2026;201:118895. 10.1016/j.trac.2026.118895.

[33] Bogusiewicz J, Burlikowska K, Łuczykowski K, Jaroch K, Birski M, Furtak J, et al. New chemical biopsy tool for spatially resolved profiling of human brain tissue in vivo. Sci Rep 2021;11:19522. 10.1038/s41598-021-98973-y.

[34] Jiang RW, Zhou W, Cypel M, Demmy TL, Shafirstein G, Garza G, et al. Minimally Invasive Chemical Biopsy Needle with Self-Wettable Extraction Phase For In Vivo Tissue Sampling During Medical Procedures. Advanced Science 2026;13:e00396. 10.1002/advs.202500396.

[35] Zhou SN, Ouyang G, Pawliszyn J. Comparison of microdialysis with solid-phase microextraction for in vitro and in vivo studies. J Chromatogr A 2008;1196–1197:46–56. 10.1016/j.chroma.2008.02.068.

[36] Cudjoe E, Bojko B, de Lannoy I, Saldivia V, Pawliszyn J. Solid-phase microextraction: a complementary in vivo sampling method to microdialysis. Angew Chem Int Ed 2013;52.

[37] Boyaci E, Lendor S, Bojko B, Reyes-Garcés N, Gómez-Ríos GA, Olkowicz M, et al. Comprehensive Investigation of Metabolic Changes Occurring in the Rat Brain Hippocampus after Fluoxetine Administration Using Two Complementary In Vivo Techniques: Solid Phase Microextraction and Microdialysis. ACS Chem Neurosci 2020;11:3749–60. 10.1021/acschemneuro.0c00274.

[38] Hassani SA, Lendor S, Neumann A, Sinha Roy K, Banaie Boroujeni K, Hoffman KL, et al. Dose-Dependent Dissociation of Pro-cognitive Effects of Donepezil on Attention and Cognitive Flexibility in Rhesus Monkeys. Biological Psychiatry Global Open Science 2021. 10.1016/J.BPSGOS.2021.11.012.

[39] Shehreen S, Hassani S-A, Lendor S, Neumann A, Sinha Roy K, Pawliszyn J, et al. Cognitive engagement induces area-specific fingerprints of dopamine, acetylcholine, serotonin, glutamate and GABA in prefrontal cortex and striatum. BioRxiv 2026:2026.05.20.726721. 10.64898/2026.05.20.726721.

[40] Gamaro GD, Xavier MH, Denardin JD, Pilger JA, Ely DR, C Ferreira MB, et al. The Effects of Acute and Repeated Restraint Stress on the Nociceptive Response in Rats. 1998.

[41] Naert G, Ixart G, Maurice T, Tapia-Arancibia L, Givalois L. Brain-derived neurotrophic factor and hypothalamic-pituitary-adrenal axis adaptation processes in a depressive-like state induced by chronic restraint stress. Molecular and Cellular Neuroscience 2011;46:55–66. 10.1016/j.mcn.2010.08.006.

[42] Parrot S, Bert L, Mouly-Badina L, Sauvinet V, Colussi-Mas J, Lambás-Señas L, et al. Microdialysis monitoring of catecholamines and excitatory amino acids in the rat and mouse brain: recent developments based on capillary electrophoresis with laser-induced fluorescence detection—a mini-review. Cell Mol Neurobiol 2003;23:793–804.

[43] Min B, Yang PS, Bohlke M, Park S, R. Vago D, Maher TJ, et al. Focused ultrasound modulates the level of cortical neurotransmitters: Potential as a new functional brain mapping technique. Int J Imaging Syst Technol 2011;21:232–40.

[44] Yang PS, Kim H, Lee W, Bohlke M, Park S, Maher TJ, et al. Transcranial focused ultrasound to the thalamus is associated with reduced extracellular GABA levels in rats. Neuropsychobiology 2012;65:153–60.

[45] Sarica C, Darmani G, Ramezanpour H, Callister M, Santyr B, Grippe T, et al. Transcranial ultrasound stimulation of motor networks in Parkinson’s disease informed by local field potential dynamics. Sci Transl Med 2026;18:eady1883. 10.1126/scitranslmed.ady1883.

[46] Lipsman N, Hynynen K, Chen R, Lozano AM. Transcranial focused ultrasound in the human brain. Neuron 2026;114:601–21. 10.1016/j.neuron.2025.11.015.

[47] Yoo S, Mittelstein DR, Hurt RC, Lacroix J, Shapiro MG. Focused ultrasound excites cortical neurons via mechanosensitive calcium accumulation and ion channel amplification. Nat Commun 2022;13:493.

[48] Oh S-J, Lee JM, Kim H-B, Lee J, Han S, Bae JY, et al. Ultrasonic Neuromodulation via Astrocytic TRPA1. Current Biology 2019;29:3386–3401.e8. 10.1016/j.cub.2019.08.021.

[49] Ozeki H, Finn IM, Schaffer ES, Miller KD, Ferster D. Inhibitory stabilization of the cortical network underlies visual surround suppression. Neuron 2009;62:578–92.

[50] Murphy BK, Miller KD. Balanced amplification: a new mechanism of selective amplification of neural activity patterns. Neuron 2009;61:635–48.

[51] Gerhard DM, Pothula S, Liu R-J, Wu M, Li X-Y, Girgenti MJ, et al. GABA interneurons are the cellular trigger for ketamine’s rapid antidepressant actions. J Clin Invest 2020;130:1336–49.

[52] Luscher B, Feng M, Jefferson SJ. Antidepressant mechanisms of ketamine: Focus on GABAergic inhibition. Adv Pharmacol 2020;89:43–78.

[53] Jena BP. Membrane fusion: role of SNAREs and calcium. Protein Pept Lett 2009;16:712– 7.

[54] Azadian MM, Kiani Shabestari S, Rajan A, Martinez PJ, Macedo N, Markarian E, et al. Clearance of intracranial debris by ultrasound reduces inflammation and improves outcomes in hemorrhagic stroke models. Nat Biotechnol 2025. 10.1038/s41587-025-02866-8.

[55] Sesack SR, Pickel VM. Prefrontal cortical efferents in the rat synapse on unlabeled neuronal targets of catecholamine terminals in the nucleus accumbens septi and on dopamine neurons in the ventral tegmental area. Journal of Comparative Neurology 1992;320:145–60.

[56] Wehr M, Zador AM. Balanced inhibition underlies tuning and sharpens spike timing in auditory cortex. Nature 2003;426:442–6.

[57] Isaacson JS, Scanziani M. How inhibition shapes cortical activity. Neuron 2011;72:231– 43.

[58] Vertes RP. A PHA-L analysis of ascending projections of the dorsal raphe nucleus in the rat. Journal of Comparative Neurology 1991;313:643–68.

[59] Bunin MA, Wightman RM. Paracrine neurotransmission in the CNS: involvement of 5-HT. Trends Neurosci 1999;22:377–82.

[60] Vázquez-Borsetti P, Cortés R, Artigas F. Pyramidal neurons in rat prefrontal cortex projecting to ventral tegmental area and dorsal raphe nucleus express 5-HT2A receptors. Cerebral Cortex 2009;19:1678–86.

[61] Nolan SO, Zachry JE, Johnson AR, Brady LJ, Siciliano CA, Calipari ES. Direct dopamine terminal regulation by local striatal microcircuitry. J Neurochem 2020;155:475–93.

[62] Schindler S, Bechtold T. Mechanistic insights into the electrochemical oxidation of dopamine by cyclic voltammetry. Journal of Electroanalytical Chemistry 2019;836:94– 101. 10.1016/j.jelechem.2019.01.069.

[63] Jiang J, Cao Y, Liu J, Zhang H, Kan G, Yu K. Mass spectrometric observation on free radicals during electrooxidation of dopamine. Anal Chim Acta 2022;1193:339403. 10.1016/j.aca.2021.339403.

[64] Paxinos G, Watson C. The rat brain in stereotaxic coordinates: hard cover edition. Elsevier; 2006.

[65] Stenroos P, Paasonen J, Salo RA, Jokivarsi K, Shatillo A, Tanila H, et al. Awake rat brain functional magnetic resonance imaging using standard radio frequency coils and a 3D printed restraint kit. Front Neurosci 2018;12. 10.3389/fnins.2018.00548.

[66] O’Reilly MA, Muller A, Hynynen K. Ultrasound Insertion Loss of Rat Parietal Bone Appears to Be Proportional to Animal Mass at Submegahertz Frequencies. Ultrasound Med Biol 2011;37:1930–7. 10.1016/j.ultrasmedbio.2011.08.001.

[67] Swanson LW. Brain maps: structure of the rat brain. Gulf Professional Publishing; 2004.

