## Supplemental Material for "Ultrasonic potentiation of ketamine neuromodulation"

[*] These authors contributed equally.

### Supplementary Figure 1

### Ultrasound transducer beam plots

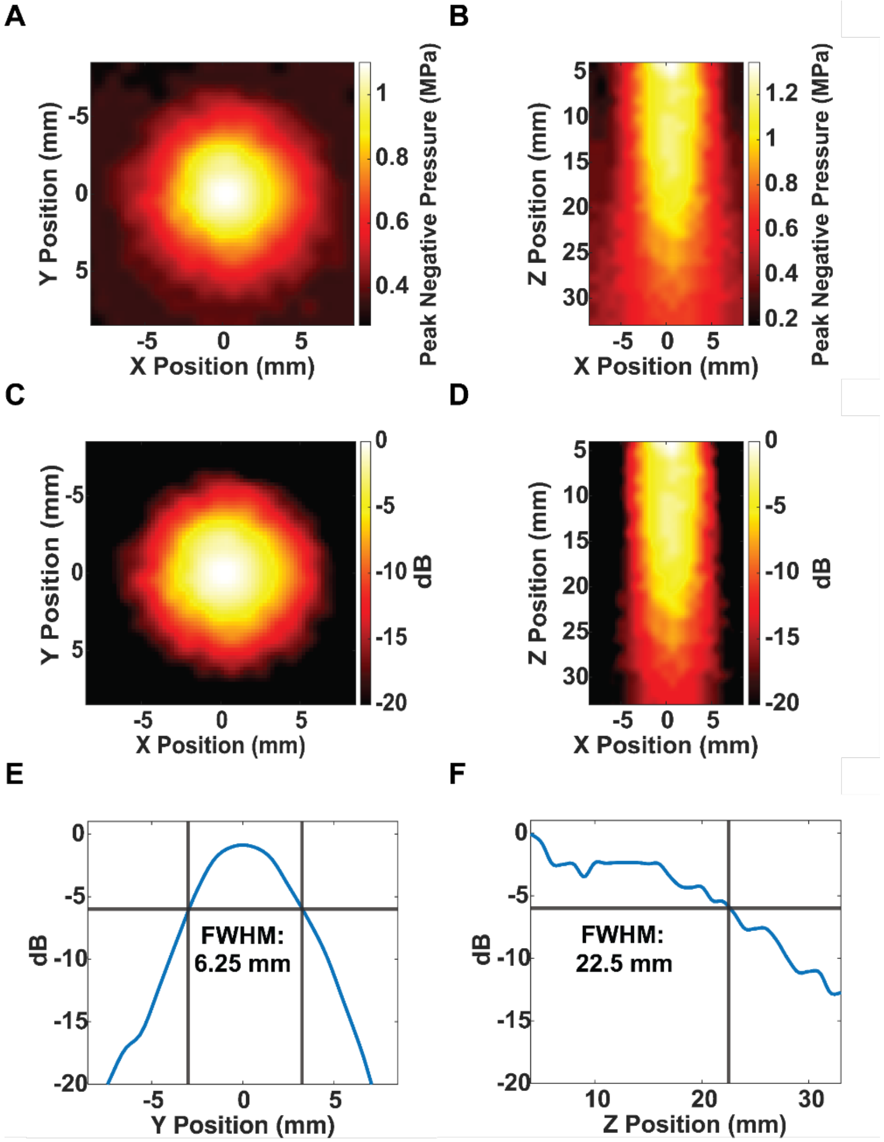

**Ultrasound transducer beam maps.** Beam maps (A, B in MPa; C, D. in dB) and 1D plots across the lateral (E) and axial (F) directions.

### Supplementary Figure 2

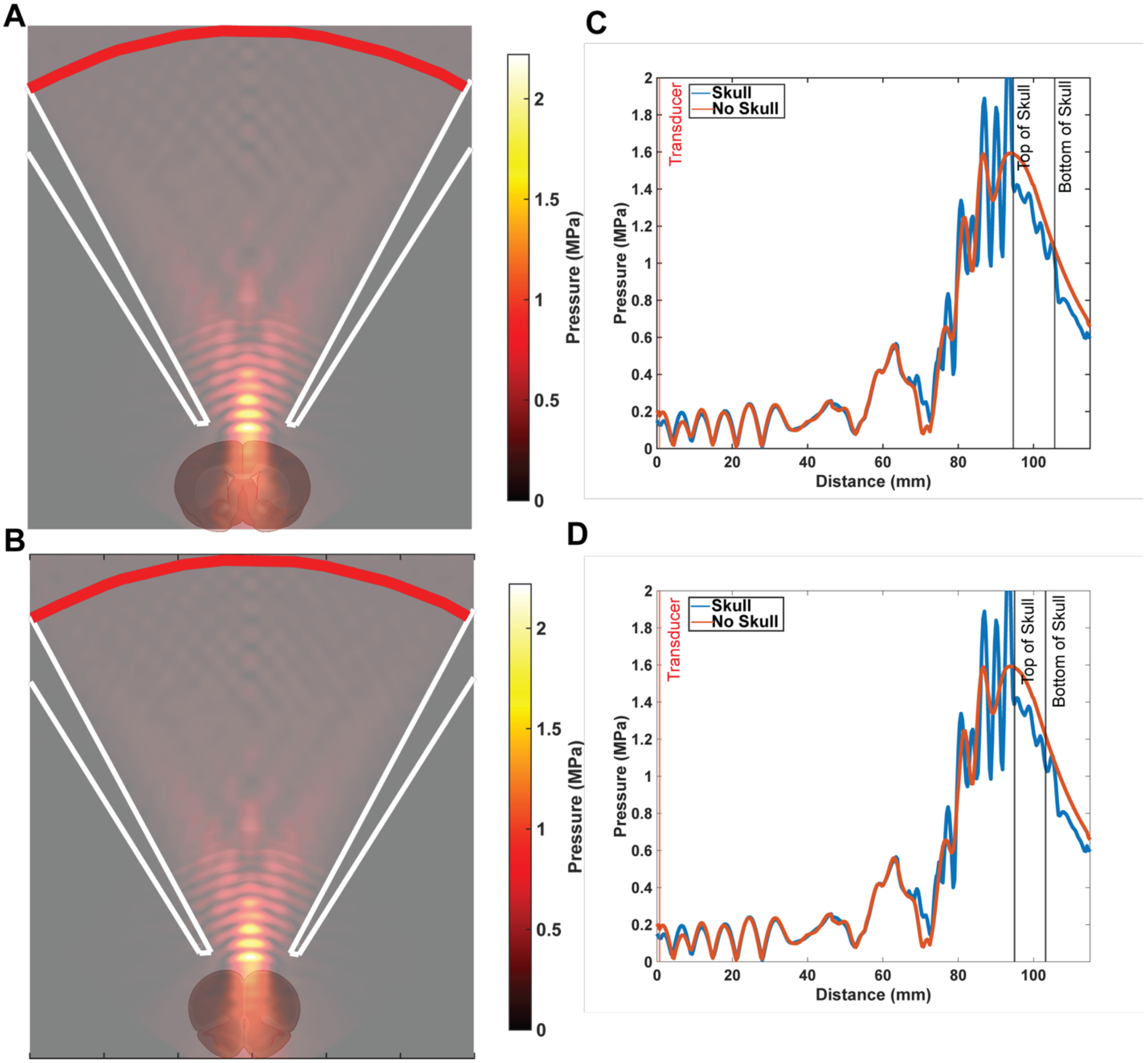

**Simulated acoustic pressure fields and axial standing wave profiles for caudal and frontolimbic TUS targets.** (A, B) Full-field pressure maps showing acoustic beam propagation through the intact rat skull. The transducer (red) is positioned above the skull surface; white lines demarcate the geometric boundaries of the acoustic coupling cone. Panels A depicts the caudal target configuration; panel B depicts the frontolimbic target configuration. (C, D) Axial pressure profiles extracted along the z-axis through the center of the transducer and rat skull–brain interface, illustrating standing wave formation proximal to the skull and across the brain volume. The blue trace represents simulations conducted with the skull present; the red trace represents free-field simulations without the skull. Panels C and D correspond to the caudal and frontolimbic configurations shown in A and B, respectively.

### Supplementary Figure 3

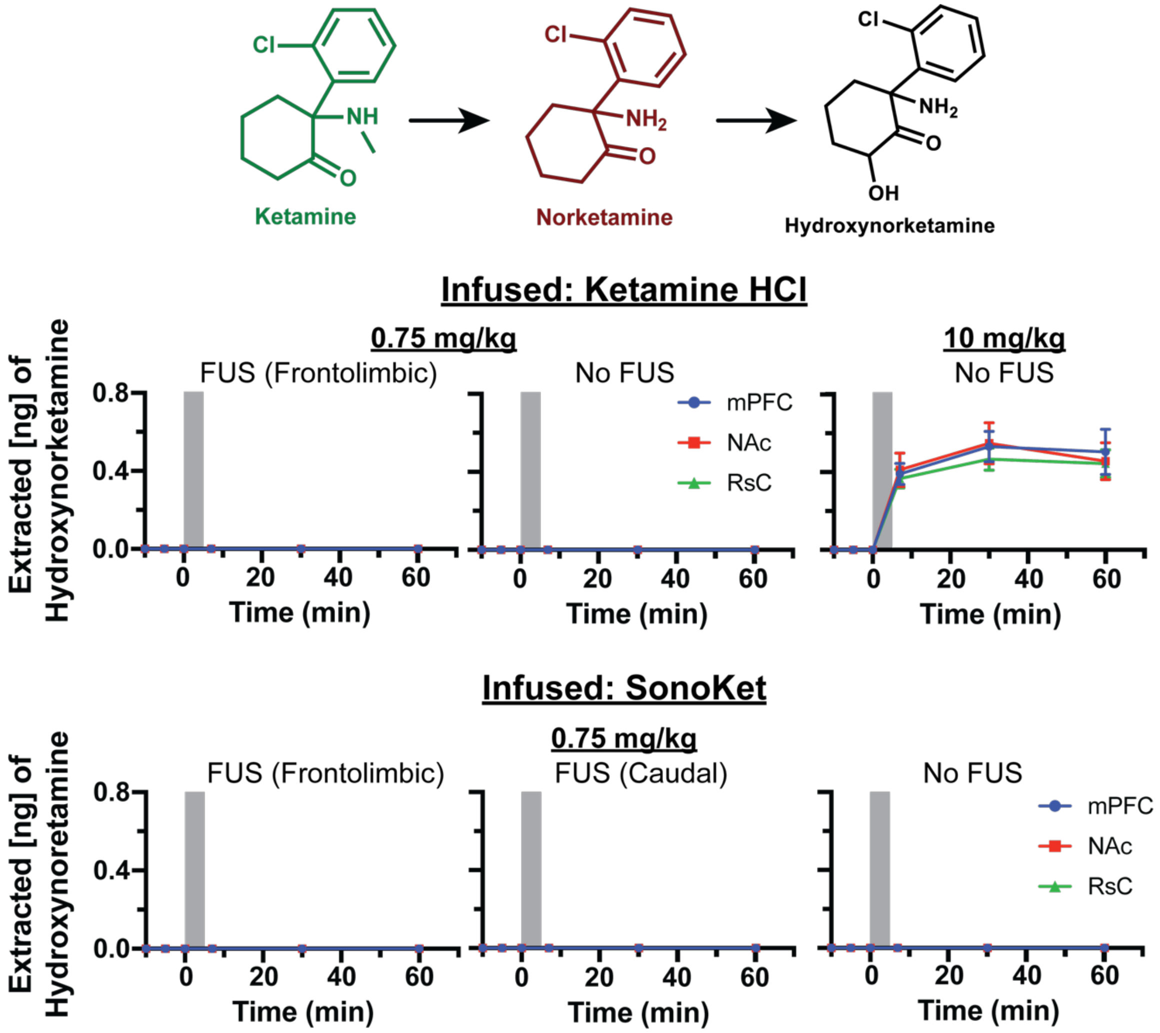

**Hydroxynorketamine profiles across mPFC, NAc, and RsC following low- and high-dose ketamine HCl or SonoKet, with and without FUS.** Hydroxynorketamine was measured by SPME sampling and coupled LC–MS/MS in the mPFC, NAc, and RsC after intravenous free ketamine HCl or SonoKet administration. Hydroxynorketamine remained near or below detectable levels after low-dose free ketamine HCl (0.75 mg $kg^{-1}$) and SonoKet (0.75 mg $kg^{-1}$), with or without FUS. In contrast, high-dose systemic ketamine HCl (10 mg $kg^{-1}$) produced measurable hydroxynorketamine across all sampled regions. Data are mean and error ± SD.; n = 4 animals per group. The grey bar indicates a 5 min duration of treatment (drug administration with or without FUS).

### Supplementary Figure 4
### (Awake restraint acclimatization)

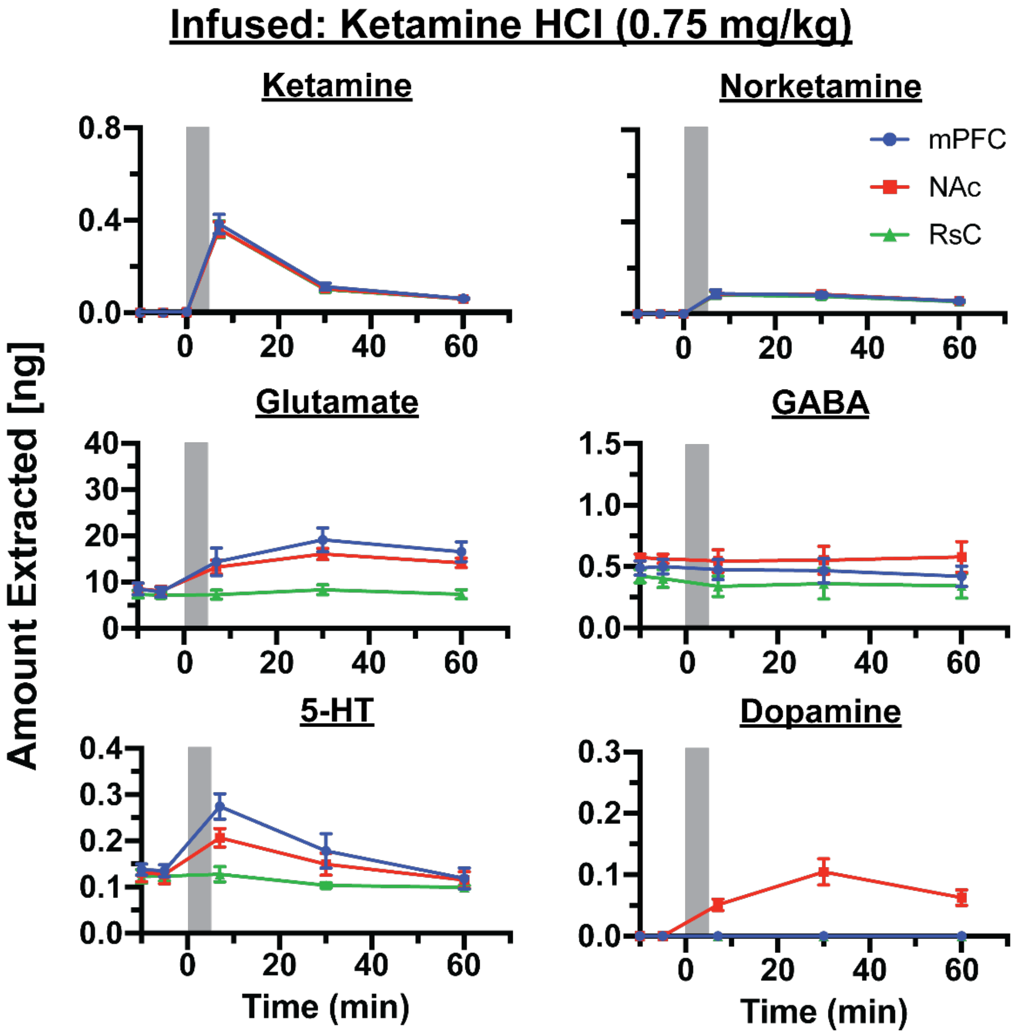

**Awake restraint-trained rats reaction to low-dose ketamine HCl with frontolimbic-targeted FUS with region-selective neurochemical responses.** Rats were acclimatized in the awake restraint bed for 90 min per day for three consecutive days before receiving low-dose free ketamine HCl (0.75 mg $kg^{-1}$) with FUS targeted to the frontolimbic region. Ketamine rapidly increased across the mPFC, NAc, and RsC and declined over time, whereas norketamine remained lower and more sustained. Glutamate and serotonin (5-HT) increased mainly in the mPFC and NAc, GABA remained relatively stable, and dopamine increased selectively in the NAc. Data are mean and error ± SD.; n = 4 animals. The grey bar indicates a 5 min duration of treatment (drug administration with or without FUS).

**Table 1: Optimized MS/MS conditions for LC-MS/MS analysis**

| Compounds | Precursor (m/z) | Product (m/z) | Dwell Time (ms) | Collision Energy (V) |
|---|---|---|---|---|
| Ketamine (q) | 238.2 | 125 | 12 | 26 |
| Ketamine | 238.2 | 220 | 12 | 10 |
| Ketamine-D4 (Internal Standard) | 242.2 | 129 | 12 | 26 |
| Norketamine (q) | 224.2 | 207 | 12 | 12 |
| Norketamine | 224.2 | 179 | 12 | 12 |
| Norketamine-D4 (Internal Standard) | 228.1 | 129 | 12 | 28 |
| Hydroxynorketamine (q) | 240.0 | 177 | 12 | 12 |
| Hydroxynorketamine | 240.0 | 195 | 12 | 12 |
| Glutamate (q) | 148.1 | 84 | 12 | 20 |
| Glutamate | 148.1 | 56 | 12 | 28 |
| Glutamate-D5 | 153.1 | 88 | 12 | 20 |
| GABA (q) | 104.1 | 86.9 | 12 | 8 |
| GABA | 104.1 | 45 | 12 | 28 |
| GABA-D6 | 110.1 | 92.9 | 12 | 8 |
| Serotonin (q) | 177.1 | 160 | 12 | 12 |
| Serotonin | 177.1 | 114.9 | 12 | 30 |
| Serotonin-D4 | 181.1 | 164 | 12 | 20 |
| Dopamine (q) | 154.1 | 91 | 12 | 21 |
| Dopamine | 154.1 | 137 | 12 | 9 |
| Dopamine-D4 | 158.1 | 140.9 | 12 | 9 |

(q) was used for quantification.

**Table 2: Optimized MS Source Parameters**

| | |
|---|---|
| Source name | AJS-ESI |
| Drying Gas Temperature (°C) | 150 °C |
| Drying Gas Flow (L/min) | 12 |
| Nebulizer Pressure (psi) | 30 |
| Sheath Gas Temperature (°C) | 350 °C |
| Sheath Gas Flow (L/min) | 11 |
| Capillary Voltage (V) | (+) 2000 |
| Nozzle Voltage (V) | 0 |
| High /Low RF voltage (v) | 80/130 |

**Table 3: Optimized Chromatographic Gradient for LC-MS/MS Analysis**

| **Time (min)** | **0** | **1** | **2.5** | **4.5** | **6** | **6.1** | **7.5** |
|---|---|---|---|---|---|---|---|
| **% B** | **2** | **2** | **20** | **100** | **100** | **2** | **2** |

**Table 4: Within-group change from baseline (paired t-test p-values)**

| | | | | | | | | | |
|---|---|---|---|---|---|---|---|---|---|
| **Glutamate** | | | | | | | | | |
| | **mPFC** | | | **NAc** | | | **RsC** | | |
| **Group** | **7 min** | **30 min** | **60 min** | **7 min** | **30 min** | **60 min** | **7 min** | **30 min** | **60 min** |
| **Saline** | <0.01 | 0.07 | 0.81 | 0.01 | 0.04 | 0.38 | 0.19 | 0.16 | 0.01 |
| **FK 0.75 FUS(Frontolimbic)** | <0.01 | <0.01 | <0.01 | <0.01 | <0.01 | <0.01 | 0.02 | <0.01 | 0.08 |
| **FK 0.75 No FUS** | <0.01 | <0.01 | <0.01 | 0.01 | <0.01 | <0.01 | 0.01 | <0.01 | 0.02 |
| **FK 10 No FUS** | <0.01 | <0.01 | <0.01 | 0.01 | <0.01 | <0.01 | 0.01 | <0.01 | 0.03 |
| **LK 0.75 FUS(Frontolimbic)** | <0.01 | <0.01 | <0.01 | <0.01 | <0.01 | 0.02 | <0.01 | <0.01 | 0.04 |
| **LK 0.75 FUS(Caudal)** | <0.01 | <0.01 | <0.01 | 0.01 | <0.01 | <0.01 | 0.01 | <0.01 | <0.01 |
| **LK 0.75 No FUS** | 0.02 | <0.01 | 0.72 | 0.02 | 0.02 | 0.06 | 0.40 | 0.03 | 0.11 |
| **GABA** | | | | | | | | | |
| | **mPFC** | | | **NAc** | | | **RsC** | | |
| **Group** | **7 min** | **30 min** | **60 min** | **7 min** | **30 min** | **60 min** | **7 min** | **30 min** | **60 min** |
| **Saline** | 0.61 | 0.15 | 0.54 | 0.66 | 0.06 | 0.06 | 0.95 | 0.44 | 0.34 |
| **FK 0.75 FUS(Frontolimbic)** | <0.01 | <0.01 | 0.76 | 0.02 | <0.01 | 0.02 | 0.07 | 0.06 | <0.01 |
| **FK 0.75 No FUS** | 0.22 | 0.03 | <0.01 | 0.03 | 0.27 | <0.01 | 0.25 | 0.06 | 0.01 |
| **FK 10 No FUS** | 0.04 | 0.62 | 0.23 | 0.05 | 0.80 | 0.04 | 0.01 | 0.32 | 0.11 |
| **LK 0.75 FUS(Frontolimbic)** | 0.52 | 0.12 | <0.01 | 0.07 | 0.03 | 0.02 | 0.65 | 0.08 | 0.09 |
| **LK 0.75 FUS(Caudal)** | 0.37 | <0.01 | 0.01 | 0.17 | 0.02 | <0.01 | 0.10 | 0.10 | 0.64 |
| **LK 0.75 No FUS** | 0.41 | 0.06 | <0.01 | 0.63 | 0.06 | 0.20 | 0.80 | 0.02 | 0.07 |
| **5HT** | | | | | | | | | |
| | **mPFC** | | | **NAc** | | | **RsC** | | |
| **Group** | **7 min** | **30 min** | **60 min** | **7 min** | **30 min** | **60 min** | **7 min** | **30 min** | **60 min** |
| **Saline** | 0.08 | 0.13 | 0.01 | 0.11 | 0.08 | 0.02 | 0.05 | 0.77 | 0.55 |
| **FK 0.75 FUS(Frontolimbic)** | <0.01 | 0.02 | 0.32 | 0.02 | 0.03 | 0.55 | 0.14 | 0.58 | 0.93 |
| **FK 0.75 No FUS** | 0.04 | 0.95 | 0.04 | <0.01 | 0.38 | 0.06 | 0.27 | 0.68 | 0.28 |
| **FK 10 No FUS** | 0.66 | <0.01 | 0.49 | 0.13 | <0.01 | 0.75 | 0.18 | 0.84 | 0.14 |
| **LK 0.75 FUS(Frontolimbic)** | <0.01 | 0.02 | 0.06 | <0.01 | 0.04 | 0.35 | 0.07 | 0.09 | 0.96 |
| **LK 0.75 FUS(Caudal)** | <0.01 | 0.54 | 0.02 | 0.78 | 0.15 | 0.23 | 0.52 | 0.13 | 0.02 |
| **LK 0.75 No FUS** | 0.04 | 0.60 | 0.02 | 0.15 | 0.51 | 0.15 | 0.25 | 0.69 | 0.79 |
| **Dopamine** | | | | | | | | | |
| | **mPFC** | | | **NAc** | | | **RsC** | | |
| **Group** | **7 min** | **30 min** | **60 min** | **7 min** | **30 min** | **60 min** | **7 min** | **30 min** | **60 min** |
| **Saline** | *n/a* | *n/a* | *n/a* | *n/a* | *n/a* | *n/a* | *n/a* | *n/a* | *n/a* |

| | | | | | | | | | |
|---|---|---|---|---|---|---|---|---|---|
| **FK 0.75 FUS(Frontolimbic)** | *n/a* | *n/a* | *n/a* | <0.01 | <0.01 | <0.01 | *n/a* | *n/a* | *n/a* |
| **FK 0.75 No FUS** | *n/a* | *n/a* | *n/a* | <0.01 | <0.01 | <0.01 | *n/a* | *n/a* | *n/a* |
| **FK 10 No FUS** | *n/a* | *n/a* | *n/a* | *n/a* | *n/a* | <0.01 | *n/a* | *n/a* | *n/a* |
| **LK 0.75 FUS(Frontolimbic)** | *n/a* | *n/a* | *n/a* | <0.01 | <0.01 | <0.01 | *n/a* | *n/a* | *n/a* |
| **LK 0.75 FUS(Caudal)** | *n/a* | *n/a* | *n/a* | *n/a* | <0.01 | <0.01 | *n/a* | *n/a* | *n/a* |
| **LK 0.75 No FUS** | *n/a* | *n/a* | *n/a* | *n/a* | *n/a* | *n/a* | *n/a* | *n/a* | *n/a* |

Paired t-test (n=4). Baseline = mean of BL1 & BL2 per replicate. Green: p<0.05; p shown to 2 dp, <0.01 below.

**Table 5: Between-group differences at each timepoint (raw p-values). Pairwise Welch t-test.**

| Glutamate | | | | | | | | | |
|---|---|---|---|---|---|---|---|---|---|
| | mPFC | | | NAc | | | RsC | | |
| **Comparison** | **7 min** | **30 min** | **60 min** | **7 min** | **30 min** | **60 min** | **7 min** | **30 min** | **60 min** |
| **Saline vs FK0.75 FUS-Frontolimbic** | 0.01 | <0.01 | <0.01 | <0.01 | <0.01 | <0.01 | <0.01 | <0.01 | <0.01 |
| **Saline vs FK0.75 NoFUS** | 0.31 | 0.01 | <0.01 | 0.63 | 0.01 | 0.03 | 0.12 | <0.01 | 0.10 |
| **Saline vs FK10 NoFUS** | <0.01 | <0.01 | <0.01 | 0.03 | <0.01 | <0.01 | 0.01 | <0.01 | 0.02 |
| **Saline vs LK0.75 FUS-Frontolimbic** | <0.01 | <0.01 | <0.01 | <0.01 | <0.01 | <0.01 | 0.02 | <0.01 | <0.01 |
| **Saline vs LK0.75 FUS-Caudal** | 0.95 | <0.01 | <0.01 | 0.22 | <0.01 | <0.01 | <0.01 | <0.01 | <0.01 |
| **Saline vs LK0.75 NoFUS** | 0.15 | 0.05 | 0.66 | 0.99 | 0.17 | 0.17 | 0.87 | 0.22 | 0.09 |
| **FK0.75 FUS- Frontolimbic vs FK0.75 NoFUS** | 0.02 | <0.01 | <0.01 | <0.01 | <0.01 | <0.01 | 0.09 | <0.01 | 0.14 |
| **FK0.75 FUS- Frontolimbic vs FK10 NoFUS** | 0.03 | 0.04 | 0.01 | 0.21 | 0.63 | 0.91 | 0.33 | 0.37 | 0.22 |
| **FK0.75 FUS- Frontolimbic vs LK0.75 FUS- Frontolimbic** | 0.18 | 0.19 | 0.05 | 0.03 | 0.23 | 0.31 | 0.41 | 0.83 | 0.41 |
| **FK0.75 FUS- Frontolimbic vs LK0.75 FUS-Caudal** | 0.01 | <0.01 | <0.01 | 0.01 | <0.01 | 0.04 | 0.02 | <0.01 | <0.01 |
| **FK0.75 FUS- Frontolimbic vs LK0.75 NoFUS** | <0.01 | <0.01 | <0.01 | <0.01 | <0.01 | <0.01 | <0.01 | <0.01 | 0.05 |
| **FK0.75 NoFUS vs FK10 NoFUS** | <0.01 | <0.01 | <0.01 | 0.03 | 0.02 | <0.01 | 0.05 | 0.03 | 0.06 |
| **FK0.75 NoFUS vs LK0.75 FUS-Frontolimbic** | <0.01 | <0.01 | <0.01 | <0.01 | <0.01 | 0.02 | 0.31 | 0.02 | 0.03 |
| **FK0.75 NoFUS vs LK0.75 FUS-Caudal** | 0.31 | 0.45 | 0.08 | 0.52 | 0.12 | <0.01 | <0.01 | <0.01 | <0.01 |
| **FK0.75 NoFUS vs LK0.75 NoFUS** | 0.06 | 0.04 | <0.01 | 0.67 | 0.13 | 0.44 | 0.13 | 0.02 | 0.63 |
| **FK10 NoFUS vs LK0.75 FUS-Frontolimbic** | 0.10 | 0.19 | 0.71 | 0.94 | 0.77 | 0.37 | 0.14 | 0.33 | 0.39 |
| **FK10 NoFUS vs LK0.75 FUS-Caudal** | <0.01 | <0.01 | <0.01 | 0.04 | 0.04 | 0.10 | 0.14 | 0.03 | <0.01 |
| **FK10 NoFUS vs LK0.75 NoFUS** | <0.01 | <0.01 | <0.01 | 0.02 | 0.01 | <0.01 | 0.02 | 0.01 | 0.05 |
| **LK0.75 FUS- Frontolimbic vs LK0.75 FUS-Caudal** | <0.01 | <0.01 | <0.01 | <0.01 | <0.01 | 0.08 | <0.01 | <0.01 | <0.01 |
| **LK0.75 FUS- Frontolimbic vs LK0.75 NoFUS** | <0.01 | <0.01 | <0.01 | <0.01 | <0.01 | 0.01 | 0.03 | <0.01 | <0.01 |
| **LK0.75 FUS-Caudal vs LK0.75 NoFUS** | 0.20 | 0.05 | <0.01 | 0.30 | 0.02 | <0.01 | <0.01 | <0.01 | <0.01 |

| GABA | | | | | | | | | |
|---|---|---|---|---|---|---|---|---|---|
| | mPFC | | | NAc | | | RsC | | |
| **Comparison** | **7 min** | **30 min** | **60 min** | **7 min** | **30 min** | **60 min** | **7 min** | **30 min** | **60 min** |
| **Saline vs FK0.75 FUS-Frontolimbic** | 0.11 | 0.14 | 0.52 | 0.13 | <0.01 | 0.01 | 0.12 | 0.05 | 0.09 |
| **Saline vs FK0.75 NoFUS** | <0.01 | 0.02 | 0.71 | 0.04 | 0.03 | 0.08 | 0.02 | 0.08 | 0.25 |
| **Saline vs FK10 NoFUS** | <0.01 | <0.01 | 0.15 | <0.01 | <0.01 | 0.28 | <0.01 | 0.02 | 0.27 |
| **Saline vs LK0.75 FUS-Frontolimbic** | 0.35 | 0.02 | <0.01 | 0.15 | 0.04 | 0.03 | 0.41 | 0.10 | 0.12 |
| **Saline vs LK0.75 FUS-Caudal** | 0.02 | <0.01 | <0.01 | 0.11 | <0.01 | <0.01 | 0.55 | 0.20 | 0.32 |
| **Saline vs LK0.75 NoFUS** | 0.83 | 0.61 | 0.50 | 0.86 | 0.83 | 0.77 | 0.69 | 0.70 | 0.91 |
| **FK0.75 FUS- Frontolimbic vs FK0.75 NoFUS** | 0.08 | 0.20 | 0.76 | 0.21 | 0.19 | 0.12 | 0.26 | 0.54 | 0.14 |
| **FK0.75 FUS- Frontolimbic vs FK10 NoFUS** | 0.01 | 0.03 | 0.23 | <0.01 | 0.80 | 0.13 | <0.01 | 0.23 | 0.21 |
| **FK0.75 FUS- Frontolimbic vs LK0.75 FUS- Frontolimbic** | 0.04 | <0.01 | <0.01 | 0.03 | 0.01 | 0.01 | 0.15 | 0.02 | 0.03 |
| **FK0.75 FUS- Frontolimbic vs LK0.75 FUS-Caudal** | 0.64 | 0.04 | 0.01 | 0.57 | 0.56 | 0.05 | 0.60 | 0.55 | 0.51 |
| **FK0.75 FUS- Frontolimbic vs LK0.75 NoFUS** | 0.20 | 0.12 | 0.19 | 0.06 | <0.01 | 0.02 | 0.14 | 0.02 | <0.01 |
| **FK0.75 NoFUS vs FK10 NoFUS** | 0.09 | 0.12 | 0.20 | 0.02 | 0.17 | 0.62 | <0.01 | 0.14 | 0.99 |
| **FK0.75 NoFUS vs LK0.75 FUS-Frontolimbic** | <0.01 | <0.01 | <0.01 | <0.01 | 0.01 | 0.01 | 0.09 | 0.03 | 0.05 |

| | | | | | | | | | |
|---|---|---|---|---|---|---|---|---|---|
| **FK0.75 NoFUS vs LK0.75 FUS-Caudal** | 0.11 | 0.23 | <0.01 | 0.62 | 0.16 | <0.01 | 0.25 | 0.79 | 0.93 |
| **FK0.75 NoFUS vs LK0.75 NoFUS** | 0.01 | 0.03 | 0.29 | 0.02 | 0.03 | 0.12 | 0.02 | 0.02 | 0.05 |
| **FK10 NoFUS vs LK0.75 FUS-Frontolimbic** | <0.01 | <0.01 | <0.01 | <0.01 | <0.01 | 0.01 | 0.01 | <0.01 | 0.04 |
| **FK10 NoFUS vs LK0.75 FUS-Caudal** | 0.02 | 0.53 | 0.09 | 0.02 | 0.68 | 0.01 | <0.01 | 0.15 | 0.93 |
| **FK10 NoFUS vs LK0.75 NoFUS** | <0.01 | <0.01 | 0.07 | <0.01 | <0.01 | 0.37 | <0.01 | 0.03 | 0.09 |
| **LK0.75 FUS- Frontolimbic vs LK0.75 FUS-Caudal** | 0.02 | <0.01 | <0.01 | 0.02 | <0.01 | <0.01 | 0.27 | 0.03 | 0.04 |
| **LK0.75 FUS- Frontolimbic vs LK0.75 NoFUS** | 0.33 | 0.05 | <0.01 | 0.16 | 0.04 | 0.02 | 0.33 | 0.07 | 0.10 |
| **LK0.75 FUS-Caudal vs LK0.75 NoFUS** | 0.08 | 0.01 | <0.01 | 0.07 | <0.01 | <0.01 | 0.70 | 0.24 | 0.27 |
| | | | | **5HT** | | | | | |
| | **mPFC** | | | **NAc** | | | **RsC** | | |
| **Comparison** | **7 min** | **30 min** | **60 min** | **7 min** | **30 min** | **60 min** | **7 min** | **30 min** | **60 min** |
| **Saline vs FK0.75 FUS-Frontolimbic** | <0.01 | <0.01 | 0.09 | 0.04 | 0.03 | 0.13 | 0.30 | 0.32 | 0.52 |
| **Saline vs FK0.75 NoFUS** | 0.02 | 0.91 | <0.01 | <0.01 | 0.17 | 0.04 | 0.38 | 0.96 | 0.28 |
| **Saline vs FK10 NoFUS** | 0.01 | <0.01 | 0.46 | <0.01 | <0.01 | 0.05 | 0.11 | 0.09 | <0.01 |
| **Saline vs LK0.75 FUS-Frontolimbic** | <0.01 | <0.01 | <0.01 | <0.01 | 0.04 | 0.08 | 0.17 | 0.06 | 0.09 |
| **Saline vs LK0.75 FUS-Caudal** | 0.02 | 0.96 | 0.05 | 0.06 | 0.41 | 0.23 | 0.10 | 0.13 | 0.17 |
| **Saline vs LK0.75 NoFUS** | 0.47 | 0.39 | 0.53 | 0.88 | 0.90 | 0.94 | 0.04 | 0.30 | 0.49 |
| **FK0.75 FUS- Frontolimbic vs FK0.75 NoFUS** | 0.19 | <0.01 | 0.01 | 0.34 | 0.01 | <0.01 | 0.11 | 0.36 | 0.21 |
| **FK0.75 FUS- Frontolimbic vs FK10 NoFUS** | <0.01 | 0.13 | 0.05 | <0.01 | 0.59 | 0.03 | 0.04 | 0.08 | 0.05 |
| **FK0.75 FUS- Frontolimbic vs LK0.75 FUS-Frontolimbic** | 0.17 | 0.47 | 0.97 | 0.16 | 0.50 | 0.24 | 0.58 | 0.19 | 0.59 |
| **FK0.75 FUS- Frontolimbic vs LK0.75 FUS-Caudal** | <0.01 | <0.01 | 0.67 | <0.01 | 0.08 | 0.91 | 0.04 | 0.38 | 0.22 |
| **FK0.75 FUS- Frontolimbic vs LK0.75 NoFUS** | <0.01 | <0.01 | 0.06 | 0.04 | 0.02 | 0.16 | 0.02 | 0.20 | 0.42 |
| **FK0.75 NoFUS vs FK10 NoFUS** | <0.01 | <0.01 | 0.03 | <0.01 | <0.01 | 0.15 | 0.22 | 0.13 | 0.23 |
| **FK0.75 NoFUS vs LK0.75 FUS-Frontolimbic** | 0.03 | <0.01 | <0.01 | 0.05 | 0.02 | 0.02 | 0.08 | 0.06 | 0.05 |
| **FK0.75 NoFUS vs LK0.75 FUS-Caudal** | <0.01 | 0.95 | <0.01 | <0.01 | 0.15 | 0.02 | 0.20 | 0.14 | 0.79 |
| **FK0.75 NoFUS vs LK0.75 NoFUS** | 0.03 | 0.57 | <0.01 | <0.01 | 0.30 | 0.03 | 0.06 | 0.49 | 0.38 |
| **FK10 NoFUS vs LK0.75 FUS-Frontolimbic** | <0.01 | 0.35 | <0.01 | <0.01 | 0.29 | 0.04 | 0.03 | 0.02 | <0.01 |
| **FK10 NoFUS vs LK0.75 FUS-Caudal** | 0.96 | <0.01 | 0.03 | 0.27 | 0.11 | 0.09 | 0.90 | 0.05 | 0.04 |
| **FK10 NoFUS vs LK0.75 NoFUS** | <0.01 | <0.01 | 0.72 | <0.01 | <0.01 | 0.14 | 0.97 | 0.23 | <0.01 |
| **LK0.75 FUS- Frontolimbic vs LK0.75 FUS-Caudal** | <0.01 | <0.01 | 0.50 | <0.01 | 0.07 | 0.31 | 0.03 | 0.71 | 0.02 |
| **LK0.75 FUS- Frontolimbic vs LK0.75 NoFUS** | <0.01 | <0.01 | <0.01 | 0.01 | 0.04 | 0.08 | 0.03 | 0.04 | 0.06 |
| **LK0.75 FUS-Caudal vs LK0.75 NoFUS** | <0.01 | 0.54 | 0.03 | 0.04 | 0.40 | 0.26 | 0.85 | 0.10 | 0.31 |
| | | | | **Dopamine** | | | | | |
| | **mPFC** | | | **NAc** | | | **RsC** | | |
| **Comparison** | **7 min** | **30 min** | **60 min** | **7 min** | **30 min** | **60 min** | **7 min** | **30 min** | **60 min** |
| **Saline vs FK0.75 FUS-Frontolimbic** | *n/a* | *n/a* | *n/a* | <0.01 | <0.01 | <0.01 | *n/a* | *n/a* | *n/a* |

| | | | | | | | | | |
|---|---|---|---|---|---|---|---|---|---|
| **Saline vs FK0.75 NoFUS** | *n/a* | *n/a* | *n/a* | <0.01 | <0.01 | <0.01 | *n/a* | *n/a* | *n/a* |
| **Saline vs FK10 NoFUS** | *n/a* | *n/a* | *n/a* | *n/a* | *n/a* | <0.01 | *n/a* | *n/a* | *n/a* |
| **Saline vs LK0.75 FUS-Frontolimbic** | *n/a* | *n/a* | *n/a* | <0.01 | <0.01 | <0.01 | *n/a* | *n/a* | *n/a* |
| **Saline vs LK0.75 FUS-Caudal** | *n/a* | *n/a* | *n/a* | *n/a* | <0.01 | <0.01 | *n/a* | *n/a* | *n/a* |
| **Saline vs LK0.75 NoFUS** | *n/a* | *n/a* | *n/a* | *n/a* | *n/a* | *n/a* | *n/a* | *n/a* | *n/a* |
| **FK0.75 FUS- Frontolimbic vs FK0.75 NoFUS** | *n/a* | *n/a* | *n/a* | 0.20 | 0.20 | 0.01 | *n/a* | *n/a* | *n/a* |
| **FK0.75 FUS- Frontolimbic vs FK10 NoFUS** | *n/a* | *n/a* | *n/a* | <0.01 | <0.01 | <0.01 | *n/a* | *n/a* | *n/a* |
| **FK0.75 FUS- Frontolimbic vs LK0.75 FUS- Frontolimbic** | *n/a* | *n/a* | *n/a* | 0.01 | <0.01 | <0.01 | *n/a* | *n/a* | *n/a* |
| **FK0.75 FUS- Frontolimbic vs LK0.75 FUS-Caudal** | *n/a* | *n/a* | *n/a* | <0.01 | <0.01 | 0.25 | *n/a* | *n/a* | *n/a* |
| **FK0.75 FUS- Frontolimbic vs LK0.75 NoFUS** | *n/a* | *n/a* | *n/a* | <0.01 | <0.01 | <0.01 | *n/a* | *n/a* | *n/a* |
| **FK0.75 NoFUS vs FK10 NoFUS** | *n/a* | *n/a* | *n/a* | <0.01 | <0.01 | 0.12 | *n/a* | *n/a* | *n/a* |
| **FK0.75 NoFUS vs LK0.75 FUS-Frontolimbic** | *n/a* | *n/a* | *n/a* | <0.01 | <0.01 | <0.01 | *n/a* | *n/a* | *n/a* |
| **FK0.75 NoFUS vs LK0.75 FUS-Caudal** | *n/a* | *n/a* | *n/a* | <0.01 | <0.01 | <0.01 | *n/a* | *n/a* | *n/a* |
| **FK0.75 NoFUS vs LK0.75 NoFUS** | *n/a* | *n/a* | *n/a* | <0.01 | <0.01 | <0.01 | *n/a* | *n/a* | *n/a* |
| **FK10 NoFUS vs LK0.75 FUS-Frontolimbic** | *n/a* | *n/a* | *n/a* | <0.01 | <0.01 | <0.01 | *n/a* | *n/a* | *n/a* |
| **FK10 NoFUS vs LK0.75 FUS-Caudal** | *n/a* | *n/a* | *n/a* | *n/a* | <0.01 | <0.01 | *n/a* | *n/a* | *n/a* |
| **FK10 NoFUS vs LK0.75 NoFUS** | *n/a* | *n/a* | *n/a* | *n/a* | *n/a* | <0.01 | *n/a* | *n/a* | *n/a* |
| **LK0.75 FUS- Frontolimbic vs LK0.75 FUS-Caudal** | *n/a* | *n/a* | *n/a* | <0.01 | <0.01 | 0.02 | *n/a* | *n/a* | *n/a* |
| **LK0.75 FUS- Frontolimbic vs LK0.75 NoFUS** | *n/a* | *n/a* | *n/a* | <0.01 | <0.01 | <0.01 | *n/a* | *n/a* | *n/a* |
| **LK0.75 FUS-Caudal vs LK0.75 NoFUS** | *n/a* | *n/a* | *n/a* | *n/a* | <0.01 | <0.01 | *n/a* | *n/a* | *n/a* |

Pairwise t-test (n=4). raw p-values. Green: significant (<0.05); p to 2 dp, <0.01 below. Each row is one group pair.

**Table 6: Between-group differences in AUC & Peak (raw p-values). Pairwise Welch t-test.**

| Glutamate | | | | | | |
|---|---|---|---|---|---|---|
| | mPFC | | NAc | | RsC | |
| **Comparison** | **AUC** | **Peak** | **AUC** | **Peak** | **AUC** | **Peak** |
| **Saline vs FK0.75 FUS- Frontolimbic** | <0.01 | <0.01 | <0.01 | <0.01 | <0.01 | <0.01 |
| **Saline vs FK0.75 NoFUS** | <0.01 | 0.03 | <0.01 | 0.06 | <0.01 | 0.01 |
| **Saline vs FK10 NoFUS** | <0.01 | <0.01 | <0.01 | 0.01 | <0.01 | 0.01 |
| **Saline vs LK0.75 FUS- Frontolimbic** | <0.01 | <0.01 | <0.01 | <0.01 | <0.01 | <0.01 |
| **Saline vs LK0.75 FUS-Caudal** | <0.01 | 0.02 | <0.01 | <0.01 | <0.01 | <0.01 |
| **Saline vs LK0.75 NoFUS** | 0.15 | 0.71 | 0.84 | 0.54 | <0.01 | 0.23 |
| **FK0.75 FUS- Frontolimbic vs FK0.75 NoFUS** | <0.01 | <0.01 | <0.01 | <0.01 | 0.31 | <0.01 |
| **FK0.75 FUS- Frontolimbic vs FK10 NoFUS** | 0.02 | 0.04 | 0.51 | 0.63 | 0.07 | 0.37 |
| **FK0.75 FUS- Frontolimbic vs LK0.75 FUS-Frontolimbic** | 0.09 | 0.19 | 0.17 | 0.18 | 0.14 | 0.83 |
| **FK0.75 FUS- Frontolimbic vs LK0.75 FUS-Caudal** | <0.01 | <0.01 | <0.01 | <0.01 | <0.01 | <0.01 |

| | | | | | | |
|---|---|---|---|---|---|---|
| **FK0.75 FUS- Frontolimbic vs LK0.75 NoFUS** | <0.01 | <0.01 | <0.01 | <0.01 | <0.01 | <0.01 |
| **FK0.75 NoFUS vs FK10 NoFUS** | <0.01 | <0.01 | 0.02 | 0.02 | 0.05 | 0.03 |
| **FK0.75 NoFUS vs LK0.75 FUS-Frontolimbic** | <0.01 | <0.01 | <0.01 | <0.01 | 0.06 | 0.02 |
| **FK0.75 NoFUS vs LK0.75 FUS-Caudal** | 0.15 | 0.52 | 0.07 | 0.03 | <0.01 | <0.01 |
| **FK0.75 NoFUS vs LK0.75 NoFUS** | <0.01 | 0.04 | <0.01 | 0.18 | <0.01 | 0.03 |
| **FK10 NoFUS vs LK0.75 FUS- Frontolimbic** | 0.19 | 0.19 | 0.62 | 0.65 | 0.24 | 0.33 |
| **FK10 NoFUS vs LK0.75 FUS-Caudal** | <0.01 | <0.01 | 0.03 | 0.05 | 0.01 | 0.02 |
| **FK10 NoFUS vs LK0.75 NoFUS** | <0.01 | <0.01 | <0.01 | 0.01 | 0.01 | 0.02 |
| **LK0.75 FUS- Frontolimbic vs LK0.75 FUS-Caudal** | <0.01 | <0.01 | <0.01 | <0.01 | <0.01 | <0.01 |
| **LK0.75 FUS- Frontolimbic vs LK0.75 NoFUS** | <0.01 | <0.01 | <0.01 | <0.01 | <0.01 | <0.01 |
| **LK0.75 FUS-Caudal vs LK0.75 NoFUS** | <0.01 | 0.03 | <0.01 | <0.01 | <0.01 | <0.01 |
| **GABA** | | | | | | |
| | **mPFC** | | **NAc** | | **RsC** | |
| **Comparison** | **AUC** | **Peak** | **AUC** | **Peak** | **AUC** | **Peak** |
| **Saline vs FK0.75 FUS-Frontolimbic** | 0.04 | 0.16 | <0.01 | 0.01 | 0.08 | 0.15 |
| **Saline vs FK0.75 NoFUS** | 0.19 | 0.18 | 0.03 | 0.02 | 0.96 | 0.08 |
| **Saline vs FK10 NoFUS** | 0.38 | 0.06 | 0.32 | 0.12 | 0.15 | 0.10 |
| **Saline vs LK0.75 FUS-Frontolimbic** | 0.06 | <0.01 | 0.05 | 0.04 | 0.23 | 0.12 |
| **Saline vs LK0.75 FUS-Caudal** | 0.04 | <0.01 | <0.01 | <0.01 | 0.25 | 0.27 |
| **Saline vs LK0.75 NoFUS** | 0.51 | 0.98 | 0.08 | 0.52 | 0.85 | 0.53 |
| **FK0.75 FUS-Frontolimbic vs FK0.75 NoFUS** | <0.01 | 0.95 | <0.01 | 0.69 | <0.01 | 0.46 |
| **FK0.75 FUS-Frontolimbic vs FK10 NoFUS** | 0.66 | 0.21 | 0.07 | 0.78 | 0.89 | 0.60 |
| **FK0.75 FUS-Frontolimbic vs LK0.75 FUS-Frontolimbic** | 0.03 | <0.01 | <0.01 | 0.02 | 0.03 | 0.05 |
| **FK0.75 FUS-Frontolimbic vs LK0.75 FUS-Caudal** | 0.55 | 0.02 | 0.58 | 0.09 | 0.14 | 0.93 |
| **FK0.75 FUS-Frontolimbic vs LK0.75 NoFUS** | <0.01 | 0.28 | <0.01 | 0.10 | 0.01 | 0.09 |
| **FK0.75 NoFUS vs FK10 NoFUS** | 0.14 | 0.24 | 1.00 | 0.61 | 0.10 | 0.93 |
| **FK0.75 NoFUS vs LK0.75 FUS-Frontolimbic** | 0.12 | <0.01 | 0.03 | 0.01 | 0.18 | 0.04 |
| **FK0.75 NoFUS vs LK0.75 FUS-Caudal** | <0.01 | 0.04 | <0.01 | 0.25 | 0.03 | 0.58 |
| **FK0.75 NoFUS vs LK0.75 NoFUS** | 0.39 | 0.28 | 0.81 | 0.08 | 0.67 | 0.05 |
| **FK10 NoFUS vs LK0.75 FUS-Frontolimbic** | 0.03 | <0.01 | 0.02 | 0.01 | 0.04 | 0.04 |
| **FK10 NoFUS vs LK0.75 FUS-Caudal** | 0.58 | 0.59 | 0.09 | 0.19 | 0.44 | 0.65 |
| **FK10 NoFUS vs LK0.75 NoFUS** | 0.24 | 0.08 | 0.91 | 0.28 | 0.12 | 0.12 |
| **LK0.75 FUS-Frontolimbic vs LK0.75 FUS-Caudal** | 0.03 | <0.01 | 0.01 | <0.01 | 0.07 | 0.05 |
| **LK0.75 FUS-Frontolimbic vs LK0.75 NoFUS** | 0.09 | <0.01 | 0.03 | 0.03 | 0.16 | 0.09 |
| **LK0.75 FUS-Caudal vs LK0.75 NoFUS** | <0.01 | 0.02 | <0.01 | 0.02 | 0.04 | 0.40 |
| **5HT** | | | | | | |

| | mPFC | | NAc | | RsC | |
|---|---|---|---|---|---|---|
| **Comparison** | **AUC** | **Peak** | **AUC** | **Peak** | **AUC** | **Peak** |
| **Saline vs FK0.75 FUS-Frontolimbic** | <0.01 | <0.01 | 0.03 | 0.03 | 0.65 | 0.19 |
| **Saline vs FK0.75 NoFUS** | 0.49 | 0.02 | 0.85 | <0.01 | 0.78 | 0.85 |
| **Saline vs FK10 NoFUS** | <0.01 | <0.01 | 0.02 | 0.08 | 0.95 | 0.09 |
| **Saline vs LK0.75 FUS-Frontolimbic** | 0.01 | <0.01 | 0.03 | <0.01 | 0.13 | 0.09 |
| **Saline vs LK0.75 FUS-Caudal** | 0.23 | 0.58 | 0.24 | 0.99 | 0.25 | 0.19 |
| **Saline vs LK0.75 NoFUS** | 0.55 | 0.47 | 0.70 | 0.88 | 0.59 | 0.33 |
| **FK0.75 FUS-Frontolimbic vs FK0.75 NoFUS** | 0.07 | 0.19 | 0.05 | 0.32 | 0.56 | 0.16 |
| **FK0.75 FUS-Frontolimbic vs FK10 NoFUS** | 0.82 | 0.31 | 0.48 | 0.11 | 0.67 | 0.04 |
| **FK0.75 FUS-Frontolimbic vs LK0.75 FUS-Frontolimbic** | 0.19 | 0.17 | 0.30 | 0.16 | 0.42 | 0.27 |
| **FK0.75 FUS-Frontolimbic vs LK0.75 FUS-Caudal** | <0.01 | <0.01 | 0.24 | 0.03 | 0.54 | 0.52 |
| **FK0.75 FUS-Frontolimbic vs LK0.75 NoFUS** | <0.01 | <0.01 | 0.02 | 0.04 | 0.46 | 0.09 |
| **FK0.75 NoFUS vs FK10 NoFUS** | 0.08 | 0.67 | 0.09 | 0.09 | 0.88 | 0.13 |
| **FK0.75 NoFUS vs LK0.75 FUS-Frontolimbic** | 0.02 | 0.03 | 0.02 | 0.05 | 0.13 | 0.08 |
| **FK0.75 NoFUS vs LK0.75 FUS-Caudal** | 0.76 | 0.02 | 0.39 | 0.03 | 0.22 | 0.18 |
| **FK0.75 NoFUS vs LK0.75 NoFUS** | 0.57 | 0.03 | 0.69 | <0.01 | 0.87 | 0.45 |
| **FK10 NoFUS vs LK0.75 FUS-Frontolimbic** | 0.15 | 0.05 | 0.14 | 0.02 | 0.21 | 0.04 |
| **FK10 NoFUS vs LK0.75 FUS-Caudal** | <0.01 | <0.01 | 0.45 | 0.15 | 0.30 | 0.08 |
| **FK10 NoFUS vs LK0.75 NoFUS** | <0.01 | 0.01 | 0.02 | 0.06 | 0.77 | 0.25 |
| **LK0.75 FUS-Frontolimbic vs LK0.75 FUS-Caudal** | 0.01 | <0.01 | 0.08 | <0.01 | 0.90 | 0.72 |
| **LK0.75 FUS-Frontolimbic vs LK0.75 NoFUS** | 0.01 | <0.01 | 0.01 | 0.01 | 0.09 | 0.06 |
| **LK0.75 FUS-Caudal vs LK0.75 NoFUS** | 0.44 | 0.95 | 0.21 | 0.90 | 0.17 | 0.13 |
| **Dopamine** | | | | | | |
| | **mPFC** | | **NAc** | | **RsC** | |
| **Comparison** | **AUC** | **Peak** | **AUC** | **Peak** | **AUC** | **Peak** |
| **Saline vs FK0.75 FUS-Frontolimbic** | *n/a* | *n/a* | <0.01 | <0.01 | *n/a* | *n/a* |
| **Saline vs FK0.75 NoFUS** | *n/a* | *n/a* | <0.01 | <0.01 | *n/a* | *n/a* |
| **Saline vs FK10 NoFUS** | *n/a* | *n/a* | <0.01 | <0.01 | *n/a* | *n/a* |
| **Saline vs LK0.75 FUS-Frontolimbic** | *n/a* | *n/a* | <0.01 | <0.01 | *n/a* | *n/a* |
| **Saline vs LK0.75 FUS-Caudal** | *n/a* | *n/a* | <0.01 | <0.01 | *n/a* | *n/a* |
| **Saline vs LK0.75 NoFUS** | *n/a* | *n/a* | *n/a* | *n/a* | *n/a* | *n/a* |
| **FK0.75 FUS-Frontolimbic vs FK0.75 NoFUS** | *n/a* | *n/a* | 0.07 | 0.20 | *n/a* | *n/a* |
| **FK0.75 FUS-Frontolimbic vs FK10 NoFUS** | *n/a* | *n/a* | <0.01 | <0.01 | *n/a* | *n/a* |
| **FK0.75 FUS-Frontolimbic vs LK0.75 FUS-Frontolimbic** | *n/a* | *n/a* | <0.01 | <0.01 | *n/a* | *n/a* |
| **FK0.75 FUS-Frontolimbic vs LK0.75 FUS-Caudal** | *n/a* | *n/a* | <0.01 | 0.04 | *n/a* | *n/a* |
| **FK0.75 FUS-Frontolimbic vs LK0.75 NoFUS** | *n/a* | *n/a* | <0.01 | <0.01 | *n/a* | *n/a* |

| | | | | | | |
|---|---|---|---|---|---|---|
| **FK0.75 NoFUS vs FK10 NoFUS** | *n/a* | *n/a* | <0.01 | <0.01 | *n/a* | *n/a* |
| **FK0.75 NoFUS vs LK0.75 FUS-Frontolimbic** | *n/a* | *n/a* | <0.01 | <0.01 | *n/a* | *n/a* |
| **FK0.75 NoFUS vs LK0.75 FUS-Caudal** | *n/a* | *n/a* | <0.01 | 0.26 | *n/a* | *n/a* |
| **FK0.75 NoFUS vs LK0.75 NoFUS** | *n/a* | *n/a* | <0.01 | <0.01 | *n/a* | *n/a* |
| **FK10 NoFUS vs LK0.75 FUS-Frontolimbic** | *n/a* | *n/a* | <0.01 | <0.01 | *n/a* | *n/a* |
| **FK10 NoFUS vs LK0.75 FUS-Caudal** | *n/a* | *n/a* | <0.01 | <0.01 | *n/a* | *n/a* |
| **FK10 NoFUS vs LK0.75 NoFUS** | *n/a* | *n/a* | <0.01 | <0.01 | *n/a* | *n/a* |
| **LK0.75 FUS-Frontolimbic vs LK0.75 FUS-Caudal** | *n/a* | *n/a* | <0.01 | <0.01 | *n/a* | *n/a* |
| **LK0.75 FUS-Frontolimbic vs LK0.75 NoFUS** | *n/a* | *n/a* | <0.01 | <0.01 | *n/a* | *n/a* |
| **LK0.75 FUS-Caudal vs LK0.75 NoFUS** | *n/a* | *n/a* | <0.01 | <0.01 | *n/a* | *n/a* |

Pairwise t-test (n=4). raw p-values. Green: significant (<0.05); p to 2 dp, <0.01 below. Each row is one group pair.
